# Single cell multiomic analyses reveal divergent effects of *DNMT3A* and *TET2* mutant clonal hematopoiesis in inflammatory response

**DOI:** 10.1101/2022.08.25.505316

**Authors:** Wazim Mohammed Ismail, Jenna A. Fernandez, Moritz Binder, Terra L. Lasho, Minsuk Kim, Susan M. Geyer, Amelia Mazzone, Christy M. Finke, Abhishek A. Mangaonkar, Jeong-Heon Lee, Liguo Wang, Kwan Hyun Kim, Vernadette A. Simon, Fariborz Rakhshan Rohakthar, Amik Munankarmy, Seul Kee Byeon, Susan M. Schwager, Jonathan J. Harrington, Melissa R. Snyder, Keith D. Robertson, Akhilesh Pandey, Eric D. Wieben, Nicholas Chia, Alexandre Gaspar-Maia, Mrinal M. Patnaik

## Abstract

*DNMT3A* and *TET2* are epigenetic regulators commonly mutated in age related clonal hematopoiesis (CH). Despite having opposed epigenetic functions, these mutations are associated with increased all-cause mortality and a low risk for progression to hematological neoplasms. While individual impacts on the epigenome have been described using different model systems, the phenotypic complexity in humans remains to be elucidated. Here we make use of a natural inflammatory response occurring during coronavirus disease 2019 (COVID-19), to understand the association of these mutations with inflammatory morbidity and mortality. We demonstrate the age-independent, negative impact of *DNMT3A* mutant CH on COVID-19-related cytokine release severity and mortality. Using single cell proteogenomics we show that *DNMT3A* mutations involve cells of myeloid and lymphoid lineages. Using single cell multiomics sequencing, we identify cell-specific gene expression changes associated with *DNMT3A* mutations, along with significant epigenomic deregulation affecting enhancer accessibility, resulting in overexpression of IL32, a proinflammatory cytokine that can result in inflammasome activation in monocytes and macrophages. Finally, we show with single cell resolution that the loss of function of DNMT3A is directly associated with increased chromatin accessibility in mutant cells. Together, these data provide a mechanistic insight into the poor inflammatory outcomes seen in *DNMT3A* mutant CH patients infected with Sars-COV2.

## INTRODUCTION

*DNMT3A* and *TET2* are key epigenetic regulator genes with opposing effects on DNA methylation. While DNMT3A is responsible for the *de novo* conversion of cytosine (C) to methylcytosine (mC), resulting in gene silencing, TET2 catalyzes the conversion of mC to 5-hydroxy-mC and subsequent oxidative metabolites, resulting in gene activation.^1^ *DNMT3A* and *TET2* are also the two most frequently mutated genes in age related clonal hematopoiesis (CH; >70%) and in spite of opposing epigenetic effects, have a convergent impact with regards to hematopoietic stem and progenitor cell (HSPC) fitness, inflammaging, low rates of progression to hematological neoplasms and increased all-cause mortality, largely due to atherosclerotic cardiovascular disease (ASCD).^2, 3, 4^ In CH, while *TET2* mutations have been associated with a myeloid lineage bias, *DNMT3A* mutations have a broader distribution, affecting myeloid and lymphoid lineage cells.^5^

Severe acute respiratory syndrome coronavirus 2 (SARS-CoV-2), the pathogen responsible for coronavirus disease 2019 (COVID-19), has resulted in an ongoing pandemic associated with significant morbidity, mortality, and long-term sequelae.^6, 7, 8^ While infections can vary from asymptomatic carrier states to severe cytokine release syndrome (CRS), acute respiratory distress (ARDS) and associated multi organ dysfunction syndrome (MODS), reasons for clinical heterogeneity have partially been investigated, with several viral (strain type) and host factors (age, comorbidities, immune status, ACE2 receptor polymorphisms, among others) congruently being involved.^9^

In a large cohort of 515 patients with COVID-19, CH was associated with severe COVID-19 outcomes, including increased mortality.^10^ However, many of these patients had underlying visceral malignancies, with several getting chemo-/ immunotherapy, and except for a non-significant trend with *PPM1D* mutations, there were no clear mutational associations detected.^10^In a subsequent study of non-cancer patients, CH was identified in 33% of 568 patients affected by COVID-19, with *DNMT3A* and *TET2* mutant (mt) CH being most frequent. In this study, neither did the presence of *DNMT3A/TET2*mt CH, nor their variant allele fractions (VAF) impact COVID-19 related outcomes^11^. Studies with smaller sample sizes and varying COVID-19 severity have demonstrated similar prevalence rates of CH, with no clear impact on outcomes^12, 13^. Autosomal mosaic chromosomal abnormalities, considered a form of CH, have been documented across large biobanks (n=768,762) and have been associated with increased infections, including a higher incidence of sepsis, respiratory tract infections, gastrointestinal and genitourinary infections.^14^

In a cohort of 243 community patients, we demonstrate an age- and comorbidity-independent adverse impact of *DNMT3A*mt CH on COVID-19 related inflammatory outcomes and over-all survival (OS). *DNMT3A*mt CH was the most frequent CH subtype, with DNA methylation studies showing a decrease in CpG site DNA methylation, mostly in distal enhancer-like elements, compared to other CH genotypes and wildtype cases. Single cell proteogenomics indicated that *DNMT3A* mutations were distributed across lymphoid and myeloid lineage cells, unlike in *TET2* mt CH, where the mutations were enriched in monocytes/macrophages. Single-cell transcriptomics demonstrated an increased expression of *IL32*, originating from NK and T lymphocytes and to some extent from classical monocytes. IL32 is a proinflammatory cytokine that can lead to monocyte/macrophage related inflammasome activation and production of cytokines such as TNF-alpha, IL8 and MIP-2.^15^. Using proximity extension assay-based proteomics (O-link), we demonstrate a negative impact of IL32 expression levels on mortality. Finally, using a combination of single cell multiome and genotyping of targeted loci with chromatin accessability (GoTChA), we identify putative epigenetic mechanisms regulating this response. These data highlight the role of *DNMT3A*mt CH in enhancing immune cell dysregulation in the context of COVID-19 infection, which may account for increased disease severity.

## RESULTS

### DNMT3Amt CH in patients with COVID-19 is associated with increased severity of cytokine release and increased age- and comorbidity-independent mortality

A cohort of 243 community-based patients with COVID-19 (alpha strain-preimmunization era) was included in the study, median age 60 years (range 19-99 years), of which 72 (29.6%) patients had evidence of CH (Supp. table 1 and **Fig. 1A**). Apart from the fact that patients with both COVID-19 and CH were older (median age for CH with COVID-19 68.5 years versus 57 years for CH without COVID-19; p < 0.0001), there were no significant differences in sex (p = 0.82), race/ethnicity (p = 0.07), hospitalization rates (p = 0.99), oxygen requirements (p = 0.79), or incidence of CRS (p=0.53). There were differences in the distribution of comorbidities between the two groups (p=0.008), with the non-CH group having a higher frequency of obesity (Supp. table 1). There, however, were no differences in baseline blood indices (mean corpuscular volume and red cell distribution width) between the two groups, in hemoglobin levels (p = 0.099), absolute neutrophil counts (ANC, p = 0.15), absolute monocyte counts (AMC; p = 0.86), or platelet counts (p = 0.41), respectively (Supp. table 2). Apart from elevated MCP-1 (monocyte chemoattractant protein-1) levels obtained at COVID-19 diagnosis in the COVID-19 CH cohort compared to the COVID-19 cohort without CH (p = 0.045), there were no other significant differences in clinically measured cytokines / chemokines, including IL1b, IL6, GM-CSF and TNF-alpha, or inflammatory surrogates like C-reactive protein (p = 0.087) and serum ferritin (p = 0.62) (Supp. table 2 and 3).

**Figure 1.**
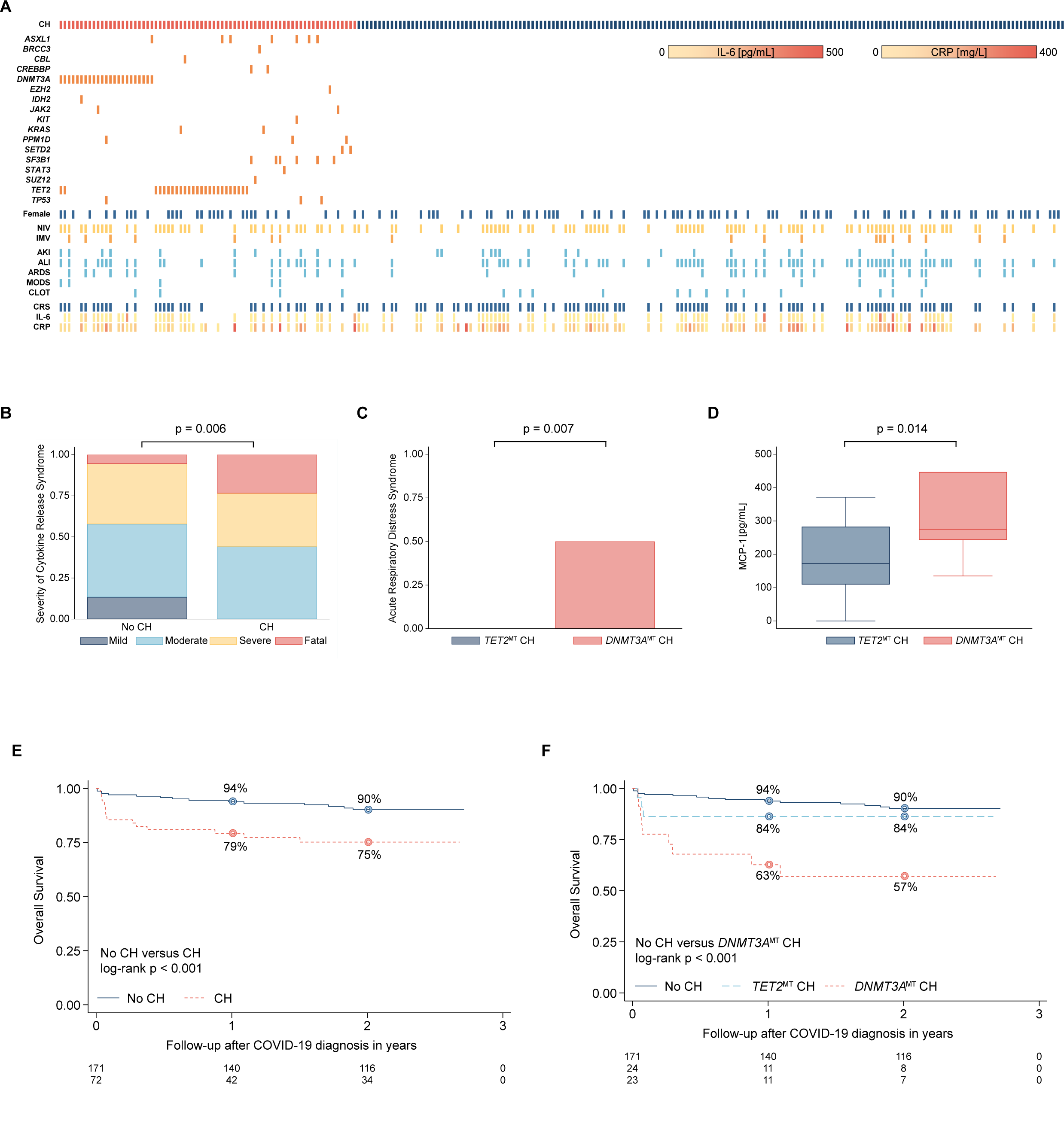
Clinical features, prevalence of clonal hematopoiesis, and disease-related outcomes of 243 patients hospitalized with COVID-19 in the pre-vaccination era. **A**, Heatmap showing the spectrum of clonal hematopoiesis (CH) mutations, sex distribution, COVID-19 related complications, prevalence of cytokine release syndrome (CRS), and serum cytokines and inflammatory markers. NIV, Non-Invasive Ventilation; IMV, Invasive Mechanical Ventilation; AKI, acute kidney injury; ALI, Acute Lung Injury; ARDS, acute respiratory distress syndrome; MODS, multiple organ dysfunction syndrome; CLOT, venous thromboembolism; CRP, C-reactive protein. **B**, Bar plots comparing the distribution of cytokine release syndrome (CRS) severity (WHO CRS severity scale) between patients with CH and without CH (No CH). There was a decrease in mild cases and an increase in fatal occurrences of CRS among patients with underlying CH (Fisher’s exact test, *p* = 0.006). **C**, Bar plots comparing the prevalence of Acute Respiratory Distress Syndrome (ARDS) among COVID-19 patients with *TET2*mt CH and *DNMT3A*mt CH. ARDS exclusively occurred in COVID-19 patients with underlying *DNMT3A*mt CH but not *TET2*mt CH (Mann-Whitney test, *p* = 0.007). **D**, Box plots comparing the serum Monocyte Chemoattractant Protein 1 (MCP-1) concentrations among COVID-19 patients with *TET2*mt CH and *DNMT3A*mt CH at the time of hospitalization. There was an increase in serum MCP-1 concentration in COVID-19 patients with underlying *DNMT3A*mt CH compared to those with *TET2*mt CH (Mann-Whitney test, *p* = 0.014). **E**, Kaplan-Meier plot showing the overall survival estimates for 243 COVID-19 patients, stratified by CH status. There was increased all-cause mortality among COVID-19 patients with underlying CH (log-rank test, *p* < 0.001). **F**, Kaplan-Meier plot showing the overall survival estimates for 218 COVID-19 patients, stratified by CH status (further stratified into *TET2*mt CH and *DNMT3A*mt CH). The increased all-cause mortality among COVID-19 patients with underlying CH was mainly driven by *DNMT3A*mt CH (log-rank test, *p* < 0.001). This association remained consistent after adjusting for age at COVID-19 diagnosis: HR 2.84 (95% CI 1.16 −6.94, *p* = 0.022).

Ninety-seven CH mutations were seen in 72 patients, with 21 (29%) having 2 CH mutations, 1 (1.3%) having 3 CH mutations and 2 (2.7%) having 4 CH mutations, respectively (**Fig. 1A**). While none of these patients underwent bone marrow (BM) biopsies, they did not have an underlying diagnosed hematological neoplasm at the time of COVID-19 detection and none of these patients demonstrated disease evolution at last follow-up (median 27 months). The most common CH mutations encountered included *DNMT3A* (n=30, 30%), *TET2* (n=26, 28%), *ASXL1* (n=7, 9.7%), *SF3B1* (n=6, 8%), *TP53* (n=3, 4%) and *PPM1D* (n=3, 4%), respectively (**Fig. 1A**). None of the *TP53* and *PPM1D* mutated patients had prior exposures to cytotoxic chemotherapy or ionizing radiation therapy. Four patients had 2 *DNMT3A* mutations, 2 patients had 2 *TET2* mutations, and one patient had both a *TET2* and a *DNMT3A* mutation. There were 4 patients with *DNMT3A* R882 hot spot mutations, commonly seen in AML and MDS. ^16^

Patients with CH and COVID-19 had higher grades of CRS in comparison to those without CH as documented by CTCAE v5.0 criteria (p = 0.006; **Fig. 1B**) and by the WHO COVID-19 severity criteria (p = 0.023).^17, 18^ There were no significant differences between the CH and no-CH groups with regards to incidence rates of acute lung injury (p = 0.8), ARDS (p = 0.49), acute kidney injury (p = 0.5), MODS (p = 0.09) and venous thromboembolism (p = 0.76) (Supp. table 4). We then compared these outcomes in the two most common CH mutant groups, *DNMT3A* and *TET2*. *DNMT3A*mt CH patients had a higher frequency of ARDS in comparison to *TET2*mt CH (p = 0.007; **Fig 1C**). The *DNMT3A*mt CH group also had significantly higher levels of MCP-1 in comparison to the *TET2*mt CH group (p = 0.014; **Fig 1D**). Both groups received similar therapeutic interventions including remdesivir, IL-6 directed monoclonal antibody therapies (tocilizumab and siltuximab), corticosteroids, and access to clinical trials (p = 0.45, Supp. table 4)). At last follow-up (median 27 months), 16 deaths (6.5%) have been documented, 10 (4%) in COVID-19 patients with CH and 6 (2.4%) in COVID-19 patients without CH.

On a univariate and multivariate survival analysis that included several clinical and laboratory variables, the presence of CH negatively impacted OS in patients with COVID-19 (p = 0.001, **Fig.1E**). The one-month estimated survival rate was 97.6% for COVID-19 patients without CH and 81.8% in COVID-19 patients with CH (median OS not reached versus 13 months).

We then analyzed the impact of *DNMT3A* and *TET2* mutations (**Supp. Fig. 1**), the two most common somatic mosaic states in our cohort on COVID-19 related morbidity and mortality. There were no significant differences between the two mutational cohorts regarding age and other comorbidities. While both *TET2*mt CH and *DNMT3A*mt CH negatively impacted survival, after adjustment for age and comorbidities, only *DNMT3A*mt CH retained an independent prognostic effect (p<0.001; **Fig. 1F**). We hence demonstrate the prevalence, clinical characteristics, and the age- and comorbidity-independent impact of *DNMT3Amt* CH on inflammatory morbidity and overall mortality, in community dwelling patients infected by the alpha strain of SARS-Cov-2.

### DNMT3Amt CH in the context of COVID-19 is associated with decreased DNA methylation at CpG residues in contrast to patients with TET2mt CH and COVID-19

Given that both *DNMT3A* and *TET2* have opposing impacts on DNA methylation, we first assessed DNA methylation status using the Illumina Infinium Methylation EPIC array on peripheral blood mononuclear cells (PBMC) from the COVID-19 and CH cohort. *DNMT3A* mutations have been associated with DNA hypomethylation at key enhancer sites in granulocytes and mononuclear cells in patients with CH, with these elements known to regulate leukocyte function, inflammation, and adaptive immune responses.^19^ We included 7 patients with CH and COVID-19 (*TET2mt* – 4 and *DNMT3Amt* – 3). Even though there were no significant global changes in DNA methylation between the two groups (p = 0.057, **Fig. 2A**), *DNMT3A*mt patients with COVID-19 demonstrated decreased methylation at highly methylated CpG sites (β > 0.75) (Kolmogorov-Smirnov p < 2.2×10^-16^; **Fig. 2B**).^20^ Site-specific differential methylation analysis also revealed an increased number of hypomethylated sites in *DNMT3A*mt patients with COVID-19 in comparison to *TET2*mt patients with COVID-19, with 10,944 hypomethylated sites and 1,160 hypermethylated sites (Δβ > 0.1 and p < 0.01; **Fig. 2C**). We then annotated the differentially methylated regions using the ENCODE Epigenomics Roadmap reference data.^21^ We found that actively transcribed states (Tx, TxWk) were more commonly hypomethylated in *DNMT3A*mt CH compared to *TET2*mt CH. Even though there were fewer hypermethylated sites, these were more common at enhancers (Enh) and promoters (TssA, TssAFlnk) (**Fig. 2D**). Pathway analysis revealed that the hypomethylated sites are in or near genes involved in many diseases and functions related to inflammation and immune response, especially in leukocyte function (**Supp. Fig. 2A, B**). Hence, we demonstrate site specific differential methylation between *DNMT3A*mt CH and *TET2*mt CH in patients with COVID-19, with more prominent hypomethylation occurring in actively transcribed regions in *DNMT3A*mt CH.

**Figure 2.**
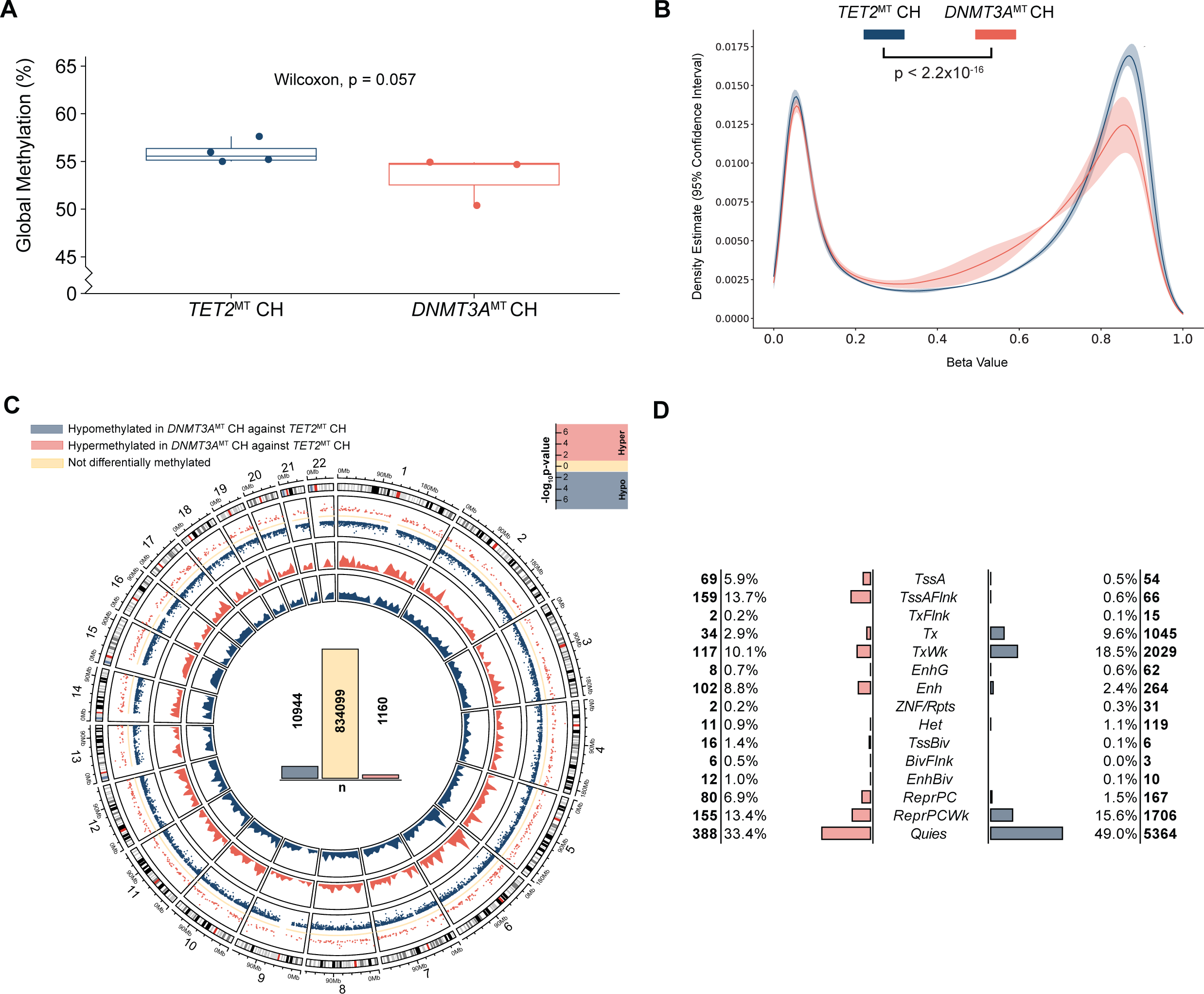
DNA methylation changes across the genome in *TET2*mt CH and *DNMT3A*mt CH patients. **A**, Boxplot showing the comparison of global DNA methylation between *DNMT3A*mt and *TET2*mt CH. There was no significant difference by Wilcoxon signed-rank test (*p* = 0.057). **B**, Density plot demonstrating DNA methylation differences between *TET2*mt CH and *DNMT3A*mt CH, primarily affecting highly methylated CpGs (β > 0.75, Kolmogorov-Smirnov test *p* < 2.2×10^-16^). **C**, Circos plot showing the number, genomic location, and density of differentially methylated regions between *TET2*mt CH and *DNMT3A*mt CH. There was an increased number of hypomethylated sites in *DNMT3A*mt CH compared to *TET2*mt CH. **D**, Functional annotation of the differentially methylated regions (with *Δβ* > 10% and *p* < 0.010) using the ENCODE Epigenomics Roadmap peripheral blood mononuclear cell reference data. Hypermethylation of enhancers (Enh) and promoters (TssA, TssAFlnk) was more commonly observed in *DNMT3A*mt CH (compared to *TET2*mt CH), whereas the hypomethylation observed in *DNMT3A* CH was predominately found at actively transcribed states (Tx, TxWk).

### Single cell proteogenomics reveal that DNMT3A mutations involve myeloid and lymphoid cell lineages, unlike TET2 mutations which bias hematopoiesis towards monocytosis

We carried out comprehensive proteogenomic assessments at single-cell resolution on PBMC, on 5 patients with COVID-19 and CH (*DNMT3A*mt--3, *TET2*mt--1, *DNMT3A* and *TET2* co-mutant--1) and 8 patients with COVID-19 and no CH (**Fig. 3A**). Given that the single cell DNA assay is an amplicon based assay and the fact that mutations in *TET2* do not have common hot spot regions, we did have *TET2*mt patients in our cohort where the mutant regions were not covered by the amplicons used in our assay, limiting the number of COVID-19+ *TET2*mt CH cases that we could genomically profile at the single cell level.^22^ Two of 3 (66%) COVID-19+ *DNMT3A*mt patients had 2 *DNMT3A* mutations each, while the COVID-19+ *TET2*mt patient also had a concomitant *CBL* mutation, with normal blood counts and no monocytosis.

**Figure 3.**
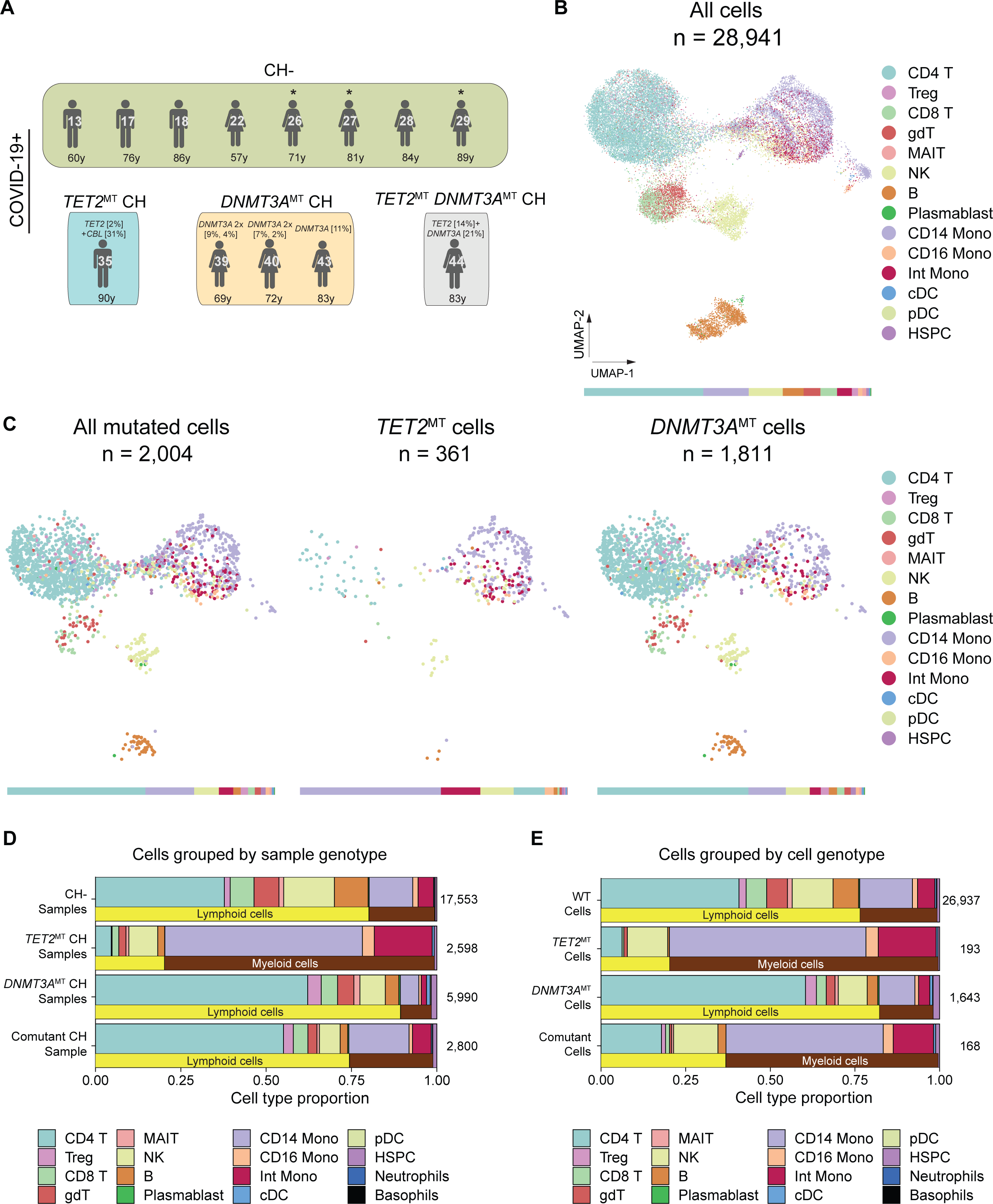
Identification of cell types carrying CH mutations by combined single-cell surface protein and genotype analysis. **A**, Overview of COVID-19 patient cohorts with *TET2*^MT^ CH, *DNMT3A*^MT^ CH, *TET2*^MT^ *DNMT3A*^MT^ (co-mutant) CH, and without CH (CH-), analyzed using single cell proteogenomics (Tapestri assay). Mutations and variant allele frequencies are shown above, and patient ages are shown below the figurines. **B**, UMAP projections showing the distribution of 28,941 cells from single cell proteogenomics analysis from 13 patient samples, colored by cell types. Key: Treg, regulatory T-cells; gdT, gamma delta T-cells; MAIT, Mucosal-associated invariant T cells; NK, natural killer cells; Mono, monocytes; Int, intermediate; cDC, classical dendritic cells; pDC, plasmacytoid dendritic cells; HSPC, hematopoietic stem and progenitor cells. The bar below shows the proportion of cells in each cell type. **C**, UMAP projections of the single cell proteogenomics data showing only mutated cells which are then stratified by *TET2* and *DNMT3A* mutated cells, demonstrating the myeloid and lymphoid lineage restriction in *TET2*mt and *DNMT3A*mt cells respectively. The bars below show the proportion of cells in each cell type. **D-E**, Bar plots showing the proportion of cells in each cell type stratifying cells by sample/patient genotype (**D**) and by cell genotype (**E**). While *TET2* mutations had a clear myeloid lineage restriction bias, *DNMT3A* mutations were seen in myeloid and lymphoid lineages, respectively.

In total, we included 28,941 single cells in the final analysis, after rigorous quality control and exclusion of cells with allele drop out, as previously described (**Fig. 3B** and methods section).^23^ Of these 28,941 sequenced cells, 2,004 (6.9%) had detectable CH mutations, of which 1,811 (90%) were *DNMT3A*mt, 361 (18%) were *TET2*mt (**Fig. 3C**) and 168 (8%) were co-mutated with both *TET2*mt and *DNMT3A*mt. In comparison to *TET2*mt CH, where CH mutations were largely present in classical and intermediate monocytes, in *DNMT3A*mt CH, the mutations were commonly seen in lymphoid lineage cells including CD4+ and CD8+ T-lymphocytes, T-regulatory cells and gamma delta T cells (**Fig. 3D-E**), a lineage bias that has previously been described.^5^

Given the smaller sample size of CH mutant cells in the COVID-19 cohort, we conducted single cell proteogenomics on an additional 4 patients with CH (**Supp.** Fig 3A) who did not have COVID-19 and cumulatively re-analyzed the data. Of 36,557 single cells successfully sequenced, 3,314 (9%) had detectable CH mutations (**Supp. Fig. 3B-C**). Among the mutated cells 1,503 (45%) were *TET2*mt, 1,643 (50%) were *DNMT3A*mt, and 168 (5%) were co-mutant cells (**Supp. Fig. 3C**). In this larger data set we once again demonstrate an enrichment of *TET2* mutations in classical and intermediate monocytes, while *DNMT3A* mutations were commonly seen in lymphoid and myeloid lineage cells, especially T-lymphocytes (**Supp. Fig. 3C-E**).

Based on these findings, we conclude that *DNMT3A* mutations are distributed in myeloid and lymphoid lineage cells, whereas *TET2* mutations have a clear myeloid (monocytic) biased distribution.

### Single cell transcriptomic analysis of DNMT3A and TET2 mutant patient samples during inflammatory response

To explore the underlying differences in expression between *DNMT3A* and *TET2* mutants in the context of SARS-CoV-2 infection we used single cell RNA-seq from PBMC from a cohort of 24 patients, which included 15 patients with COVID-19 and no CH, 6 COVID-19 patients with *TET2* mutations and 3 COVID-19 patients with *DNMT3A* mutations (**Fig. 4A**). By pooling 78,083 cells from all 24 patients and subjecting them to dimensionality reduction using PCA and UMAP, followed by cell type identification using SingleR (see methods section and **Supp. Fig. 4A**), we identified the typical repertoire of lymphoid and myeloid cells (**Fig. 4B-C**). For the comparison of the different cell types, we also included published data from healthy individuals separated into two age groups (under and over 50 years).^24^ The most noticeable difference was the enrichment of classical and intermediate monocytes in patients with *TET2* mutations, while patients with *DNMT3A* mutations show enrichment of CD8+ and gamma delta T lymphocytes, plasma blasts, NK cells and non-classical monocytes (**Fig 4 C** and **Supp. Fig. 4 B**).

**Figure 4.**
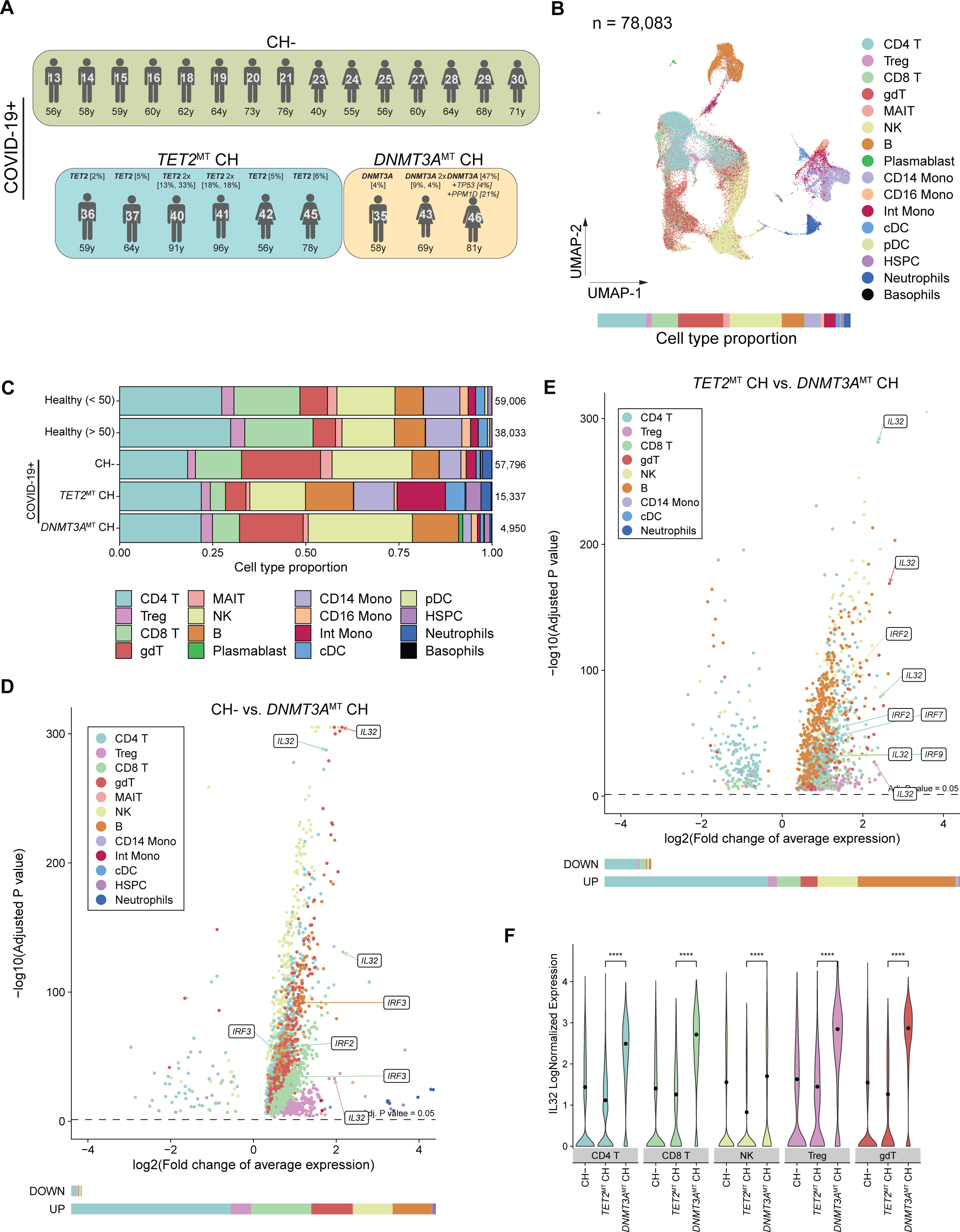
Identification of biomarkers of COVID-19 severity associated with CH mutations. **A**, Overview of COVID-19 patient cohorts without CH (CH-), *TET2*^MT^ CH and *DNMT3A*^MT^ CH, analyzed using scRNA-seq. Mutations and variant allele frequencies are shown above, and patient ages are shown below the figurines. **B**, UMAP projections showing the distribution of 78,083 cells from single cell RNA-seq analysis from 24 patient samples, colored by cell types identified using SingleR. Key: Treg, regulatory T-cells; gdT, gamma delta T-cells; MAIT, Mucosal-associated invariant T cells; NK, natural killer cells; Mono, monocytes; Int, intermediate; cDC, classical dendritic cells; pDC, plasmacytoid dendritic cells; HSPC, hematopoietic stem and progenitor cells. The bar below shows the proportion of cells in each cell type. **C**, Proportion of cells in each cell type stratified by 5 conditions as shown in y-axis, in the scRNA-seq analysis. The healthy cohort (from Stephenson et al. PMID: 33879890) is further stratified by age: below 50 and above 50 years, respectively. **D-E**, Volcano plots showing significantly differentially expressed genes (adjusted *p* < 0.05, Wilcoxon rank sum test) in comparisons between CH- and *DNMT3A*^MT^ CH, (**D**) and between *TET2*^MT^ CH and *DNMT3A*^MT^ CH (**E**), both in the context of COVID-19. Cells from each cell type were tested independently. Bars below the volcano plots show the proportion of genes per cell type that are down- and up-regulated in the *DNMT3A*^MT^ CH in each comparison. **F**, Violin plots showing expression of *IL32* in cell types where *DNMT3A*^MT^ CH patients had significantly higher expression compared to *TET2*^MT^ CH. Black dots show mean expression. ****: *p* <= 0.0001 (Wilcoxon rank sum test).

To further investigate a potential association of *DNMT3A* mutations with COVID-19 severity, we performed differential gene expression analysis within each cell type, first comparing COVID-19+ *DNMT3A*mt CH cells with COVID-19+ no CH cells and found an overall increase in *IL32* expression in CD4+ and CD8+ T lymphocytes, regulatory T cells and NK cells, a potential biomarker of severity that has not been documented in plasma samples from patients with COVID-19.^25^ (**Fig 4D**). We then compared COVID-19+ *DNMT3A*mt CH cells with COVID-19+ *TET2*mt CH cells and found the same cell type specific overexpression pattern for *IL32*, as described above (**Fig. 4E, F**). Pathway analysis of differentially expressed genes in this comparison revealed an enrichment of genes involved in lymphocyte proliferation, migration of blood cells, cytotoxicity of lymphocytes and NK cells and joint inflammation (**Supp. Fig. 4C**).

Given that IL32 has not specifically been implicated in COVID-19 severity, we analyzed the abundance of cytokines in patients with COVID-19 from our cohort of 223 assessable patients using a multiplex proteogenomic panel (O-link-methods section for details). While there were no significant differences in IL32 levels between *DNMT3A*mt and *TET2*mt CH COVID-19 patients, or between COVID-19 patients with and without CH, when we assessed this cohort for COVID-19 related morbidity and mortality by looking at relative levels of IL32, we found that COVID-19 patients with higher IL32 levels had a higher mortality, in comparison to those with lower levels (**Supp. Fig. 4D**).

Here we demonstrate a lymphoid lineage enrichment for *DNMT3A*mt CH in comparison to *TET2*mt CH, along with an inflammatory transcriptional signature resulting in overexpression of *IL32* in *DNMT3A*mt CH, with higher IL32 protein levels correlating with increased mortality in the context of COVID-19.

### Epigenetic up-regulation of IL32 occurs due to increased chromatin accessibility of a transcriptional program seen in CH patients with DNMT3A mutations

Since *DNMT3A* and *TET2* are known to regulate chromatin accessibility with opposing effects across the genome, we conducted single-cell profiling of both gene expression (scRNA-seq) and open chromatin (scATAC-seq) from the same PBMC samples using the 10X Genomics Multiome platform (methods for details).^26^ From a cohort of 11 COVID-19 patients, which included 6 patients without CH, 3 patients with *TET2*mt CH and 2 patients with *DNMT3A*mt CH were selected (**Supp. Fig. 5A**). We pooled 25,725 single cells and performed dimensionality reduction using PCA and UMAP analysis, followed by cell type identification by mapping the expression data from Multiome onto the scRNA-seq data and transferring labels using Azimuth (**Supp. Fig. 5C**). From each cell type, we were able to gather expression signatures and open chromatin profiles, expressed in cut site counts (number of sites cut by the Tn5 transposase – a direct measure of accessibility of chromatin to the transposase). Using this technology, we were able to validate the lymphoid lineage enrichment in *DNMT3A*mt CH, comprising of CD4+ T-lymphocytes, regulatory T cells, B-lymphocytes, and plasma blasts, in comparison to *TET2*mt CH, where monocytic enrichment was more prominent (classical and intermediate monocytes, dendritic cells and CD8+ T-lymphocytes; **Fig 5A**, **Supp. Fig. 5B**). As chromatin accessibility is reflective of the active enhancer and promoter structure and is strongly associated with DNA methylation status, we aimed to provide mechanistic insights on the deregulation of the epigenetic landscape in *DNMT3A*mt CH in comparison with *TET2*mt CH. Notably, analysis of global distribution of cut sites and differentially accessible peaks showed that there was increased chromatin accessibility in *DNMT3A*mt CH, especially in the two main cell types overexpressing *IL32*, CD4+ T lymphocytes and NK cells (**Fig. 5B, Supp. Fig. 5D**). Moreover, co-accessibility analysis revealed an enrichment of *cis*-regulatory interactions associated with expression of up-regulated genes such as *IL32* in *DNMT3A*mt CH cells by identifying several candidate enhancers linked to the transcription start site of *IL32* (**Fig. 5C**).^27^ The epigenomic landscape around *IL32* containing the ENCODE *cis*-regulatory elements mapped with cell specific open chromatin regions identified by scATAC analysis also allowed us to identify overlapping hypomethylated CpG regions in *DNMT3A*mt CH patients (cg01100763, cg09294055, cg04519177) obtained from bulk PBMC DNA methylation data, suggesting a direct link between loss of methylation and increased chromatin accessibility. (**Fig. 5C,D**). To understand transcription factors (TF) likely to be affected by *DNMT3A* and *TET2* CH mutations, we first performed analysis of differentially accessible TF binding sites by differential enrichment analysis directly from ChIP-seq datasets found in the literature.^28^ Expression of some of these transcription factors was also analyzed in the same cell types to ensure that the chromatin accessibility analysis could also reflect changes in expression levels (**Supp Fig. 5E, F**). Interestingly, the IRF family of transcription factors was enriched both in transcription (scRNA) and in TF activity (scATAC) in *DNMT3A*mt CH, suggesting that a specific transcriptional program driven by IRF is involved in the pro-inflammatory response, particularly in CD4+ T lymphocytes. Finally, to assess if the epigenetic dysregulation in *DNMT3A*mt CH can be traced back to the mutant cells, we performed genotyping of targeted loci with single-cell chromatin accessibility (GoTChA) in two patient samples with high VAF for a known *DNMT3A* loss of function mutation (**Supp. Fig. 6-8**).^29^ Comparing number of cutsites in wild type and mutant cells from the same sample revealed a similar pattern of increased open chromatin in *DNMT3A*mt CH cells both globally (**Fig. 5E**) and in a genomic locus specific manner, at CpG sites affected around the *IL32* locus (**Fig. 5F**). With one exception (Locus A in patient 2), *DNMT3A* mutations were directly associated with higher ATAC signal, indicating a direct link between loss of function of the DNA methyltransferase activity and increased chromatin accessibility.

**Figure 5.**
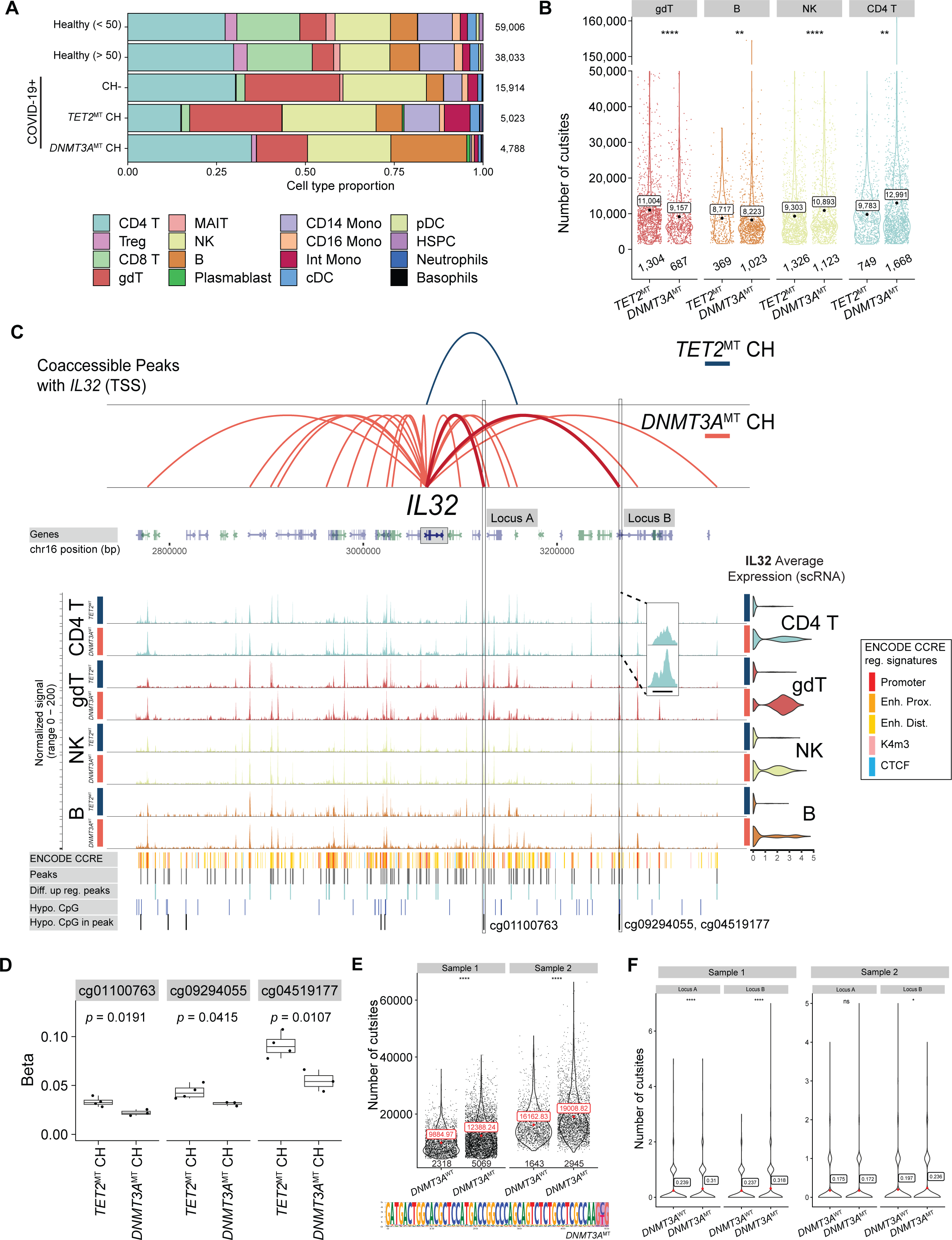
Characterization of epigenetic deregulation in CH patients with *DNMT3A* mutations. **A**, Proportion of each cell type identified in the scRNA-seq data from the 10X Multiome platform stratified by 5 conditions as shown in y-axis. The healthy cohort is further stratified by age: below 50 and above 50 years, respectively. Key: Treg, regulatory T-cells; gdT, gamma delta T-cells; MAIT, Mucosal-associated invariant T cells; NK, natural killer cells; Mono, monocytes; Int, intermediate; cDC, classical dendritic cells; pDC, plasmacytoid dendritic cells; HSPC, hematopoietic stem and progenitor cells. **B**, Violin plots showing cell type specific changes in chromatin accessibility measured as the total number of cut sites (sum of TF-IDF normalized cut site counts across all peaks; scATAC-seq data from Multiome) in each cell type. Only cell types with more than 100 cells are shown. Black dots show the mean value. ns: *p* > 0.05, *: *p* <= 0.05, **: *p* <= 0.01, ***: *p* <= 0.001, ****: *p* <= 0.0001 (Wilcoxon rank sum test). NK and CD4 T cells showed significant increase in chromatin accessibility in *DNMT3A*^MT^ CH patients compared to *TET2*^MT^ CH in the context of COVID-19. **C**, Coverage plot showing epigenomic dysregulation of *IL32* in *DNMT3A*^MT^ CH compared to *TET2*^MT^ CH, through multiomics analysis. The plot shows the co-accessible peaks with *IL32* transcription start site (TSS) in *TET2*^MT^ CH patients and *DNMT3A*^MT^ CH patients, both with COVID-19 (co-accessibility score > 0.1; blue and red arcs), the chromatin accessibility signal per group of cells, *IL32* gene expression (violin plots; right), candidate *cis*-regulatory elements predicted by ENCODE (colored-coded bars), open-chromatin peaks (grey bars), differentially accessible peaks that are more accessible in *DNMT3A*^MT^ CH patients compared to *TET2*^MT^ CH patients in CD4 T cells (light blue bars), CpG sites hypomethylated in *DNMT3A*^MT^ CH patients compared to *TET2*^MT^ CH patients (dark blue bars) and CpG sites overlapping open-chromatin regions (black bars) around the *IL32* gene locus. Labeled loci A and B are chr16:3123999-3124965 and chr16:3263558-3264913, respectively. These loci are regions where *DNMT3A*^MT^ CH patients gained accessibility in CD4 T cells, overlapped with hypomethylated CpG sites and gained co-accessibility with *IL32* transcription start site (TSS). **D**, Box plots showing methylation levels (β values) per patient in *TET2*^MT^ CH and *DNMT3A*^MT^ CH cohorts (both with COVID-19) at three hypomethylated CpG sites shown in panel C. The middle line represents the median, the lower and upper edges of the rectangle represent the first and third quartiles and the lower and upper whiskers represent the interquartile range (IQR) × 1.5. The groups were compared using Wilcoxon rank sum test. **E-F**, Violin plots showing significant increase in chromatin accessibility in *DNMT3A*^MT^ cells compared to *DNMT3A*^WT^ cells as determined by GoTChA analysis. The data shown are the total number of cut sites (**E**) and the number of cut sites at loci A and B from panel C (**F**), in *DNMT3A* wild-type and *DNMT3A* mutant cells from two samples (*DNMT3A*^MT^clonal cytopenias of undetermined significance) profiled using GoTChA. Red dots show the mean value. Mutation site in the *DNMT3A* gene is shown in the bottom of panel E. ns: *p* > 0.05, *: *p* <= 0.05, **: *p* <= 0.01, ***: *p* <= 0.001, ****: *p* <= 0.0001 (Wilcoxon rank sum test).

## DISCUSSION

Clonal hematopoiesis is defined by the acquisition of somatic mutations in HSPC, with the capacity to expand over time, with evolving cell intrinsic and extrinsic selection pressures.^2, 3, 4^ CH is ubiquitous in the aging population, with studies demonstrating hematopoiesis to be largely oligoclonal in the elderly.^30^ *DNMT3A* and *TET2* are the two most common age-related CH mutated genes, with both being critical regulators of DNA methylation.^2, 3, 4, 31^ While age related CH is associated with a low risk of hematological neoplasms, its presence is associated with increased all-cause mortality, largely due to cardiovascular disease.^2, 3, 4, 31, 32^ This is believed to be secondary to pervasive inflammatory transcriptional priming and inflammasome activation associated with these mutations.^1, 33, 34^ While the clinical impact of these two mutations is convergent, their impact on the epigenome is not. *DNMT3A* mutations are mostly loss of function mutations that lead to protein instability and loss of methyltransferase activity, resulting in DNA hypomethylation, whereas *TET2* mutations are either truncating or hypomorphic, abrogating the catalytic activity of TET2, resulting in DNA hypermethylation.^19, 33, 35, 36^ In addition, lineage restriction analysis has shown that while *DNMT3A* mutations involve myeloid and lymphoid cell lineages, *TET2* mutations are more myeloid restricted with a clear myelomonocytic bias.^5, 37^

In this study, using COVID-19 as a model for severe inflammation, using bulk and single cell multiomics on patient samples obtained prior to the advent of the SARS-CoV-2 vaccine, we demonstrate the negative impact of CH on inflammatory morbidity (CRS) and mortality. We show that while this was accounted for by increasing age in *TET2*mt CH patients; *DNMT3A*mt CH remained an independent adverse prognosticator. While several host and viral factors impacting COVID-19 severity have been described, we demonstrate the relevance of *DNMT3A*mt CH in this context.^6, 7, 8, 10, 38^ Prior studies have shown a conflicting impact of CH on COVID-19 related morbidity and mortality.^10, 11, 12, 13^ While there could be several confounding factors explaining these discrepancies, our study population included unselected community dwelling individuals infected by the alpha strain of SARS-CoV-2, prior to immunization and without any underlying hematological or visceral neoplasms, or immunodeficiency states.

The frequency of CH in the COVID-19 cohort was 29.6%, with *DNMT3A* (30%) and *TET2* (28%) being the two most mutated CH genes, consistent with prior observations.^2, 3^ While the *DNMT3A* and *TET2*mt CH groups were well balanced with regards to baseline blood counts and comorbidities, *DNMT3A*mt CH patients had higher levels of MCP-1, had a higher likelihood of ARDS and had higher grades of CRS as judged with CTCAE v5.0 criteria and by the WHO COVID-19 severity criteria.^17, 18^ There was no further stratification of this effect based on CH mutational VAF or the number of CH mutations. None of the patients included in this cohort, at last follow-up, had evidence of a hematological neoplasm. MCP-1, also called CCL2, is a key chemokine that regulates the migration of monocytes and macrophages in response to inflammation and has been implicated as a biomarker of COVID-19 severity in the recent past.^39,40^

We then conducted methylation studies using the Illumina Methylation EPIC array in select cases, and while there were no global differences in DNA methylation, site-specific analysis revealed an increased number of hypomethylated sites in *DNMT3A*mt versus *TET2*mt patients with COVID-19. Using the ENCODE Epigenomics Roadmap reference data,^21^ we demonstrate that actively transcribed states (Tx, TxWk) were more commonly hypomethylated in *DNMT3A*mt CH compared to *TET2*mt CH, with pathway analysis revealing that the hypomethylated sites were in or near genes involved in several diseases and functions related to inflammation. This is consistent with prior observations assessing DNA methylation on whole blood samples in patients with CH, CH associated cytopenias and AML.^19^ Given the lack of a scalable single cell methylation assay, we were not able to validate these findings at the single cell level, appropriately adjusting for somatic mosaicism and duly acknowledge this limitation.

We then assessed the distribution of *DNMT3A* and *TET2*mt CH in the COVID-19 cohort using single-cell proteogenomics and validate observations from prior lineage restriction analyses that while *DNMT3A*mt CH involved myeloid and lymphoid lineage cells, *TET2*mt CH had a clear myeloid restriction, with a myelomonocytic bias.^5^ These observations were also validated with single cell RNA and multiome-seq data. Enrichment of *TET2* mutations were in classical and intermediate monocytes, reflective of a granulocyte monocyte-biased hematopoiesis (GMP-bias) and classical monopoiesis, which has been well documented in *TET2*-driven hematological neoplasms such as chronic myelomonocytic leukemia and might explain the differential impact seen on inflammatory morbidity and mortality seen in the context of COVID-19.^41, 42, 43^ Differential gene expression analysis comparing *DNMT3A*mt CH patients with those without CH, and those with *TET2*mt CH, demonstrated an overall increase in *IL32* expression in CD4+ and CD8+ T lymphocytes, regulatory T cells and NK cells in patients with *DNMT3A*mt CH versus those without CH and those with *TET2*mt CH. Pathway analysis of differentially expressed genes revealed an enrichment of genes involved in lymphocyte proliferation, migration of blood cells, cytotoxicity of lymphocytes and NK cells and joint inflammation. On O-link based cytokine analysis, while there were no significant differences in IL32 levels between *DNMT3A*mt and *TET2*mt CH COVID-19 patients, or between COVID-19 patients with and without CH, relative increments in IL32 levels were associated with a higher mortality, in comparison to those with lower levels.

IL32 is a proinflammatory cytokine, initially detected in activated NK cells and T-lymphocytes, whose expression is strongly enhanced by microbes, mitogens, and inflammatory stimuli.^44^ It can amplify production of other inflammatory cytokines including IL1b, IL6 and TNF-a and has not been reported as a biomarker of severity in COVID-19, or CH, thus far. We speculate that *IL32* expression in *DNMT3A*mt CH is enhanced in the context of inflammatory stimuli such as COVID-19. To better understand the regulatory mechanism behind *IL32* overexpression in *DNMT3A*mt CH, we conducted single cell multiome profiling using the 10X Genomics Multiome platform. On a global distribution analysis of cut sites and differentially accessible peaks, we found increased chromatin accessibility in *DNMT3A*mt CH, especially in CD4+ T lymphocytes and NK cells, the two cell types with predominant *IL32* overexpression. Co-accessibility analysis revealed an enrichment of *cis*-regulatory interactions associated with expression of *IL32* in *DNMT3A*mt CH cells, identifying candidate enhancers linked to the transcription start site of *IL32*. We found that the IRF family of transcription factors was enriched both in transcription and in TF activity in *DNMT3A*mt CH, particularly in CD4+ T lymphocytes. Finally, to address issues with somatic mosaicism, we performed genotyping of targeted loci with single-cell chromatin accessibility (GoTChA) in two patients with *DNMT3A*mt clonal cytopenia’s and found a similar pattern of increased open chromatin in *DNMT3A*mt CH cells both, globally and in a genomic locus specific manner, at CpG sites affected around the *IL32* locus.

In summary, our results validate *DNMT3A*mt CH as an age and comorbidity independent risk factor for severe COVID-19. *DNMT3A*mt CH involves myeloid and lymphoid lineage cells and in the context of inflammatory stimuli, results in the overexpression of *IL32*, a highly proinflammatory cytokine, predominantly in T and NK cells. This regulation in part is mediated by *DNMT3A*mt associated changes in chromatin accessibility, allowing for transcription factors like the IRF family of TF, to mediate transcriptional activation at *IL32* promotor sites.

## ACKNOWLEDGMENTS

The authors thank the Genome Analysis Core and the Biospecimens Accessioning and Processing (Mayo Clinic) for technical support, and Alessandro Gardini for help with access to their published datasets. This work was supported by the Mayo Clinic Center for Individualized Medicine, by the Mayo Clinic Center for Biomedical Discovery to W.M.I. and A.G.M. and the DOD Ovarian Cancer Research Program (W81XWH2110475 to A.G.M.). MMP would also like to acknowledge the NCI for R01 grant R01CA272496.

## AUTHOR CONTRIBUTIONS

M.P., A.G.M., N.C., conceived and designed the study with the help of M.B., T.L., W.M.I., J.F., and J.J.H. J.F., T.L., A.Mazzone, C.M.F., K.H.K., V.A.S., F.R.R., A.Munankarmy, S.K.B., M.R.S. and J-H.L. performed experiments. W.M.I., T.L., M.B., J.F. M.K., S.M.G., A.A.M., S.M.S. and L.W. analyzed the data. M.P., A.G.M. and M.B., wrote the manuscript with the help of W.M.I., J.F., K.R., N.C., A.P., and E.W. All authors critically revised and approved the final version of the manuscript.

## COMPETING INTERESTS

M.M.P., has received research funding from Kura Oncology, Epigenetix, Solutherapeutics, Polaris and StemLine Pharmaceuticals.

## MATERIALS AND METHODS

### Authorizations, patient cohorts, cell collection and sorting

This study was conducted at the Mayo Clinic in Rochester, Minnesota, after approval from the Mayo Clinic Institutional Review Board (IRB-20-005400). In all cases, diagnosis was according to the 2016 iteration of the WHO classification of myeloid malignancies.^45^ PB and BM samples were collected in EDTA tubes after informed consent.

### Target Capture Chip Assay for Bulk Sequencing

Sample DNA was extracted from peripheral blood mononuclear cells isolated by gradient centrifugation and re-suspended in a concentration of 500 ng in 50µl of low TE buffer. Paired-end indexed libraries were prepared using the Sureselect XT Low Input Library prep protocol on the Agilent Bravo liquid handler following the manufacturer’s protocol (NewEngland Biolabs, Ipswitch, MA, and Agilent Technologies, Ankeny, IA). Briefly, 200ng of target DNA was fragmented using the Covaris LE220 plus sonicator. The settings of duty factor 30%, peak incident power (PIP) 450, cycles per burst 200, time 180 seconds, generated double-stranded DNA fragments with blunt or sticky ends with a fragment size mode of between 150-200bp. The ends were repaired using the Sureselect End-Repair-A-Tailing enzyme mix. Adapter ligated DNA fragments were size-selected to enrich for 200 bp inserts (∼320 bp total library size) using AMPURE XP bead purification. The size-selected adapter-modified fragments were enriched, and specific indexes were added by 12 cycles of PCR using universal index primers. The concentration and size distribution of the libraries was determined on an Agilent Bioanalyzer DNA 1000 chip.

The Custom Capture hybrid-target enrichment probes were designed using Agilent SureSelect design software (Agilent Technologies, Santa Clara, CA). The targeted gene panel was comprised of 62,962 single probes with size 1.805Mbp, and covered the coding regions, UTRs, and overlapping intron/exon regions for 205 genes described and / or enriched for CH mutations (see Appendix: Target Gene Panel). The custom capture was carried out using the Agilent Bravo liquid handler following Agilent’s SureSelect XT Low. Purified capture products were then amplified using the SureSelect Post-Capture primer mix for 14 cycles. Libraries were validated and quantified on the Agilent Bioanalyzer. Samples were sequenced by 150 paired end reads, 21 samples to a Flow Cell, on an Illumina NovaSeq 6000 SP (Illumina, SanDiego, CA).

Secondary bioinformatics analysis included quality assessment and alignment to the hg19 build reference genome using Novoalign (Novocraft Technologies, Malaysia), followed by GATK based single nucleotide and small insertion / deletion variant calling, structural variation discovery, and annotation. The quality of sequencing chemistry was evaluated using FastQC. After alignment, PCR duplication rates and percent reads mapped on target were used to assess the quality of the sample preparations. Realignment and recalibration steps were implemented in the GATK.^46^ Somatic single nucleotide variations (SNVs) were then genotyped using SomaticSniper, whereas insertions and deletions were called by GATK Somatic Indel Detector.^46, 47^ Each variant in coding regions was functionally annotated by SnpEff, SAVANT, ClinVar, dbNSFP, OMIM, and the Human Gene Annotation Database to predict biological effects.^48, 49, 50, 51, 52, 53^ Each variant was also annotated with allele frequency from the Exoma Aggregation Consortium.^54^ Variants of significant interest were visually inspected using IGV.^55^ The total list of all variants was internally compared for internal duplicates indicative of false positives, and variants of concern are additionally hand annotated for identification using Alamut Software. Interpretation for relevant alterations included absence in international normal variant allele databases (GnomAD, ExAC), deleterious effect on protein function by multiple phenotype prediction models, somatic and functional annotation in literature, consequence of variant (nonsense, truncating, etc.) and location proximal to important domains.

### Single-Cell Proteogenomics

Single-cell DNAseq + proteogenomics was performed using the Mission Bio Tapestri platform according to the manufacturer’s specifications. Briefly, a cocktail of pre-titered oligo-linked antibodies targeting 42 unique cell-surface markers and 3 antibodies for isotype control, TotalSeq^TM^ D Human Heme Oncology Kit (BioLegend®, San Diego, CA [https://www.biolegend.com/en-us/totalseq/single-cell-dna]) was utilized to capture the cell immunophenotype, along with targeted myeloid scDNAseq genotyping panel targeted against 45 genes with 312 amplicons covering relevant genes dysregulated in myeloid disorders (designed and manufactured by Mission Bio, San Francisco, CA [https://designer.missionbio.com/catalogpanels/Myeloid]). See Appendices: Myeloid Panel Coverage and Myeloid Panel Design for specific genomic coordinates. Cryopreserved PBMC patient samples were gently thawed, washed, and run through a Dead Cell Removal Kit (Miltenyi, Auburn, CA). The resulting viable cell fraction was counted, blocked with Human TruStain FcX^TM^ (BioLegend®) and re-suspended at a concentration of 25,000 cells/µL. This fraction was incubated with BioLegend TotalSeq Kit Cocktail described, washed, and filtered through a Flowmi cell strainer (MilliporeSigma, St. Louis, MO), and quantified using a Countess II cell counter. The cells were then diluted to a concentration of 4,000 cells per μL in Cell Buffer. Next, 35 μL of cell suspension was loaded onto a microfluidics cartridge and cells were encapsulated on the Tapestri instrument followed by cell lysis with protease digestion followed by heat inactivation using a thermal cycler. The cell lysate was reintroduced into the cartridge and processed such that each cell possessed a unique molecular barcode. Amplification of the targeted DNA regions was performed by incubating the barcoded DNA emulsions in a thermocycler as follows: 98 °C for 6 min (4 °C per s); ten cycles of 95°C for 30s, 72°C for 10s, 61°C for 9 min, 72°C for 20s (1 °C per s); ten cycles of 95°C for 30s, 72°C for 10s, 48°C for 9 min, 72°C for 20s (1°C per s); and 72°C for 6 min (4°C per s). Emulsions were broken, DNA digested and purified with 0.72X AMPure XP beads (Beckman Coulter). The beads were pelleted and washed with 80% ethanol and the generated by amplifying DNA libraries with Mission Bio V2 index primers in the thermocycler using the following program: 95°C for 3 min; 10 cycles of 98°C for 20s for DNA libraries and 16-20 cycles for protein libraries, 62°C for 20s, 72°C for 45s; 72°C for 2 min. Final libraries were purified with 0.69X AMPure XP beads. All libraries were sized and quantified using an Agilent Bioanalyzer and pooled for sequencing on an Illumina NovaSeq 6000 SP with 2 x 150bp multiplexed runs. FASTQ files generated by sequencers were processed using the Tapestri Pipeline V2 which handles adapter trimming, sequence alignment (BWA), barcode correction, cell finding and variant calling (using GATK 4.1.7 haplotype caller). Generated loom and H5 files were then processed with Tapestri Insights v2.2 (Mission Bio) and/or the Python-based Mosaic package (GitHub). Tapestri Insights analysis used default filter criteria (for example, genotype quality ≥30 and reads per cell per target ≥10) or whitelisting of known variants and annotation-based information (including ClinVar and DANN). Only cells with complete genotype information for all variants (previously detected in bulk sample) were included for downstream processing.

### Cell type identification for the proteogenomics platform

A rigorous quality control was performed on antibody-derived tag (ADT) count profiles of the Tapestri single-cell proteogenomic data. We first filtered out cells with less than 200 or more than 50,000 total ADT counts, and then removed cells with less than 10 unique measured ADTs. After ADT-based filtering, one sample was excluded from the analysis as there were less than 500 cells remaining for the sample. ADT counts for each surface protein were then scaled using counts per 10 million with a pseudocount of +1 and normalized using the centered-log ratio (CLR) transformation. Subsequently, following step II of the “dsb” (denoised and scaled by background) protocol, normalized ADT profiles of antibodies for isotype control were used to remove cell-to-cell technical noise.^56^ In result, we obtained a normalized and denoised ADT count matrix representing 42 different surface protein expression profiles of 36,557 single cells from 17 patients. This full matrix was used to generate UMAP plots following dimensionality reduction using PCA.

To identify cell types for cells profiled by the Tapestri platform, we compared ADT count profiles of the Tapestri data with that of a reference CITE-seq data, i.e., Azimuth reference.^57^ Among the 42 cell-surface markers targeted by our Tapestri analysis, 36 were also targeted by the reference CITE-seq analysis. In the CITE-seq analysis, expression of 8 out of the 36 shared markers were assayed using two different antibodies targeting the same antigen (CD3, CD4, CD11b, CD38, CD44, CD45, CD56, and CD138). For these antigens, we used geometric means of ADT counts for both antibodies. CD5, CD7, CD10, CD33, CD62L, and FceRIa were the markers that were targeted by the Tapestri platform but not by the reference CITE-seq analysis. ADT count matrices of the 36 shared markers for the Tapestri and CITE-Seq analyses were scaled, normalized, and denoised as described above (for denoising CITE-seq data, 3 IgG antibodies were selected as isotype controls). Subsequently, for each cell from the Tapestri data, 20 nearest neighbor cells from the CITE-seq data were identified using Euclidian distance, and cell type was called based on their identities using majority vote.

### Single-cell profiling of PBMCs (single-cell RNA-seq and 10X Genomics Multiome)

Frozen PBMCs were thawed in a 37°C water bath for 3-5 min until no ice was visible. Cells were washed twice with 1 mL PBS + 0.04% BSA and pelleted (300×g for 5 min at 4°C). Dead cells were removed according to the Demonstrated Protocol: Removal of Dead Cells from Single Cell Suspensions for Single Cell RNA Sequencing (10X Genomics). Using the MACS Dead Cell Removal Kit (Miltenyi Biotec), the pellet was resuspended in 100 μL Dead Cell Removal MicroBeads and incubated for 15 minutes at room temperature. After incubation, the cell suspension was diluted with 1X Binding Buffer and applied to an MS column. The dead cells were retained on the column and the live cells passed through the column and were collected. After dead cell removal, the samples were washed twice with 1 mL PBS + 0.04% BSA and the cell concentration was determined using a Cellometer K2 cell counter (Nexcelom Biosciences). The cells were then aliquoted for scRNA-seq and Multiome (see Methods section).

### Single-cell RNA-seq

The cells were first counted and measured for viability using either the Vi-Cell XR Cell Viability Analyzer (Beckman-Coulter, Brea, CA) or a basic hemocytometer and light microscope. The barcoded Gel Beads were thawed from −80°C and the cDNA master mix was prepared according to the manufacturer’s instruction for Chromium Next GEM Single Cell 3’ Library and Gel Bead Kit (10x Genomics, Pleasanton, CA). Based on the desired number of cells to be captured for each sample, a volume of live cells was mixed with the cDNA master mix. A per sample concentration of 400,000 cells per milliliter or better is required for the standard targeted cell recovery of approximately 4000 cells. The stock concentration requirements would not change for higher cell recovery numbers. The cell suspension and master mix, thawed Gel Beads and partitioning oil were added to a Chromium Single Cell G chip. The filled chip was loaded into the Chromium Controller, where each sample was processed and the individual cells within the sample were captured into uniquely labeled GEMs (Gel Beads-In-Emulsion). The GEMs were collected from the chip and taken to the bench for reverse transcription, GEM dissolution, and cDNA clean-up. The resulting cDNA contained a pool of uniquely barcoded molecules. A portion of the cleaned and measured pooled cDNA continued to library construction, where standard Illumina sequencing primers and a 10x Genomics unique i7 Sample index were added to each cDNA pool. All cDNA pools and resulting libraries were measured using Qubit High Sensitivity assays (Thermo Fisher Scientific, Waltham, MA) and Agilent Bioanalyzer High Sensitivity chips (Agilent, Santa Clara, CA).

Libraries were sequenced at between 40,000 and 50,000 fragment reads per cell following Illumina’s standard protocol using the Illumina cBot and HiSeq 3000/4000 PE Cluster Kit (Illumina, San Diego, CA). The flow cells were sequenced as 100 × 2 paired end reads on an Illumina HiSeq 4000 HD using HiSeq 3000 / 4000 sequencing kit and HCS v3.4.0.38 collection software. Base-calling was performed using Illumina’s RTA version 2.7.7.

### Single-cell Multiome

After approximately 4,000 cells were aliquoted for scRNA-seq, the remainder were used for single-cell Multiome ATAC + Gene Expression (10X Genomics). Nuclei were isolated according to the Demonstrated Protocol: Nuclei Isolation for Single Cell Multiome ATAC + Gene Expression (10x Genomics, CG000365 Rev A). Briefly, cells were added to a 2.0 mL low binding tube and centrifuged (300×g for 5 min at 4°C) using a swinging bucket rotor. The supernatant was removed, and the cell pellet was resuspended in 100 µL of chilled 10x Genomics Lysis Buffer (10 mM Tris-HCl pH 7.4, 10 mM NaCl, 3 mM MgCl_2_, 0.1% Tween-20, 0.1 % NP-40 Substitute, 0.01% digitonin, 1% BSA, 1 mM DTT, 1 U/μL RNase inhibitor 40 U/mL) by pipette-mixing 10 times. Cells were incubated on ice for 3 min, followed by dilution with 1 mL of chilled Wash Buffer (10 mM Tris-HCl pH 7.4, 10 mM NaCl, 3 mM MgCl_2_, 0.1% Tween-20, 1% BSA, 1 mM DTT, 1 U/mL RNase inhibitor 40 U/mL). Nuclei were then centrifuged (500×g for 3 min at 4°C), and the supernatant was slowly removed. The nuclei were washed one additional time with 1 mL Wash Buffer. Nuclei were resuspended in chilled diluted nuclei buffer (1X Nuclei Buffer (10x Genomics), 1 mM DTT, 1 U/mL RNase inhibitor 40 U/mL); the concentration was determined using a Cellometer K2 cell counter (Nexcelom Biosciences) and the samples were adjusted to a concentration appropriate for our targeted nuclei recovery. The single-cell ATAC library construction and gene expression library construction were carried out as described in the Chromium Next GEM Single Cell Multiome ATAC + Gene Expression User Guide (CG000338 Rev A). ATAC and GEX libraries were sequenced separately on an HiSeq 4000 (Illumina) before demultiplexing, alignment to the reference genome, and post-alignment quality control.

### DNA Methylation

Genomic DNA was isolated and checked for quality by standard protocols. 1 µg genomic DNA then underwent bisulfite treatment using the TrueMethyl oxBS Module (Tecan Genomics, Männedorf, Switzerland) according to the manufacturer’s specifications. Briefly, DNA was purified using magnetic beads, denatured, and underwent bisulfite conversion followed by desulfonation and purification. The TrueMethyl converted DNA samples were then eluted in 10 µL and then processed through the Illumina Infinium MethylationEPIC BeadChip array (Illumina, San Diego, CA) protocol. In brief, 7 µL of converted DNA was denatured with 1 µL of 0.4N sodium hydroxide prior to whole genome amplification on the MSA4 plate. All other steps were followed as per the manufacturer’s guidelines.

Quality control of Infinium MethylationEPIC BeadChips was performed via the Genome Studio Methylation Module (Illumina). Subset-quantile Within Array Normalization (SWAN) was performed on the Infinium MethylationEPIC BeadChip IDAT files via the R package “minfi”.^58, 59^ The resultant β-values are used to identify changes in DNA methylation (Δβ) between groups. Unless otherwise noted, a change in absolute methylation level of 10% (Δβ > |0.1|) and a *p* value of < 0.01 were considered significant.

CpG site relation within chromatin states was annotated using Bedtools v2.27.1 to the genome annotations provided for PBMCs by the Roadmap Epigenomics project.^60^ The functional annotation of differentially methylated CpGs located within gene bodies and promoters was generated through the use of QIAGEN IPA (QIAGEN Inc., https://digitalinsights.qiagen.com/IPA). Gene ontology associated with the differentially methylated CpGs within non-coding regions was performed using GREAT.^61^

### Genotyping of Targeted loci with single-cell Chromatin Accessibility (GoT-ChA)

The assay was performed following the published protocol (Myers et al., 2022) with some modifications. 10,000 nuclei were captured for each sample. The following primers were utilized to specifically amplify genotyping fragment (DNMT3A R882). GoT-ChA R1N-F, 5’-TCGTCGGCAGCGTCAGATGTGTATAAGAGACAGGATGACTGGCACGCTCCAT-3’; GoT-ChA-R, 5’-CTAAGCAGGCGTCAGAGGAG-3’; GoT-ChA_nested-R, 5’-BiosG/CCTTGGCACCCGAGAATTCCATCCTGCTGTGTGGTTAGACG-3’. The underlined sequences are locus-specific. After index PCR, DNAs were digested with AatII and MscI to monitor the specificity of amplified DNAs. Final libraries were quantified using a Qubit dsDNA HS Assay Kit (Thermo Fisher Scientific, #Q32854) and were analyzed by the Fragment Analyzer (Advanced Analytical Technologies; AATI; Ankeny, IA) using the High Sensitivity NGS Fragment Analysis Kit (Cat. #DNF-486). The libraries were sequenced on a NovaSeq 6000 at the Mayo Clinic Genome Analysis Core. ATAC libraries were sequenced to a depth of 25,000 read pairs per nucleus and GoT-ChA libraries were sequenced to 5,000 read pairs per nucleus.

### Olink

We used Olink Explore 1536 panel assay (Olink Proteomics [Uppsala, Sweden]), which uses proximity extension assay technology coupled to a readout methodology based on next generation sequencing (Illumina NovaSeq 6000, NextSeq 550, and NextSeq 2000; all manufactured by Illumina; appendix 1 p 2), to quantify protein targets, as previously described.^62^

### Statistical analysis

Distribution of continuous variables was statistically compared using Mann-Whitney or Kruskal-Wallis tests, while nominal or categorical variables were compared using the Chi-Square or Fischer’s exact test. Time to event analyses used the method of Kaplan-Meier for univariate comparisons using the log-rank test. OS was calculated from the date of diagnosis to date of death or last follow-up, while AML-free survival (LFS) was calculated from date of diagnosis to date of AML transformation or death. Poisson regression models were fit to compare the numbers of different cell types across groups. These Poisson models incorporated an offset term to reflect the total number of cells for a given patient (to be able to compare cell types across consistent intervals in the setting of varying total cell counts: ln(*Y*) = ln(*N*) + *β*_0_ + *β*_1_*X*_1_ + *ε*).

### Single-cell data analysis

Single-cell RNA-seq: Sequenced reads from the droplet libraries were processed using 10x Genomics Cell Ranger v6.0.1.^63^ The reads were aligned to the pre-built human reference transcriptome GRCh38 -v2020-A (July 7, 2020) provided by 10X Genomics. Read trimming, alignment, UMI counting, and cell calling were performed by Cell Ranger. Doublet prediction on scRNA-seq data was done using Scrublet v0.2.1 with default parameters.^64^ Downstream processing was done using Seurat v4.0.4.^65^ Count matrices from all samples were combined and batch-corrected using Seurat v4 integration method. Count matrix from each sample was log-normalized, scaled to mean 0 and variance 1, and dimensionality reduced using PCA on the top 2000 variable genes across all samples. The reciprocal PCA and reference-based integration options were applied in the anchor finding step due to large data size. Four patient samples, one from each sex and each condition (COVID-19^+^ / CH^-^ and COVID-19^+^ / CH^+^), were chosen as references, and PCs 1-50 were used for the reference-based integration. The integrated data included a total of 85,019 cells from 24 patient samples. Cells with more than 50% of reads mapped to mitochondrial genes, those with less than 200 unique genes detected and those that were predicted as doublets by Scrublet were removed. After QC filtering the number of cells was reduced to 78,083. Genes that were not expressed in at least 3 cells were also removed from downstream analysis. Uniform manifold approximation and projection (UMAP) was made using the top 50 PCs obtained by running PCA on the integrated (batch-corrected) gene expression matrix. Cell type identification was done with SingleR v1.10.0 using immune data from celldex v1.6.0 as reference.^66, 67^

Single-cell Multiome: Sequenced reads from the gene expression (GEX) and DNA accessibility (ATAC) droplet libraries of the Multiome assay were processed using 10x Genomics Cell Ranger ARC v2.0.0. The reads were aligned to the pre-built human reference genome GRCh38 -v2020-A-2.0.0 (May 3, 2021) provided by 10X Genomics. Read trimming, alignment, duplicate marking (ATAC), UMI counting (GEX), peak calling (ATAC) and joint cell calling were performed by Cell Ranger. Downstream processing was done using Seurat v4.0.4 and Signac v1.4.0.^68^ GEX and ATAC count matrices were integrated (batch-corrected) independently using Seurat. Sample GEX count matrices were integrated following the same steps as used for the scRNA-seq data. Default options (CCA and pairwise anchor-finding) were used in the integration anchor finding step, since the data size was smaller than our scRNA-seq data. To merge the ATAC data from all samples, a common peak set was created by merging peaks from all samples using the reduce function from the R package GenomicRanges. Peaks that were smaller than 20 base pairs or larger than 10000 base pairs after merging were removed. The count matrix for each sample with the new common peak set was then recalculated using Signac. The count matrices were normalized using Term Frequency -Inverse Document Frequency (TF-IDF) and dimensionality reduced using singular value decomposition (SVD) using only peaks with non-zero counts in at least 20 cells, these two steps together known as latent semantic indexing (LSI) generating LSI components. The samples were then integrated using Seurat V4 integration with the reciprocal LSI (rLSI) method used on LSI components 2 to 50 (since the first LSI component correlates with sequencing depth) in the pairwise anchor finding step. The UMAP was calculated using the integrated LSI components 2 to 50. Seurat’s weighted nearest neighbor (WNN) algorithm was used on principal components 1 to 50 (GEX) and integrated LSI components 2 to 50 (ATAC) together to obtain a combined UMAP projection of both GEX and ATAC counterparts of the complete scMultiome dataset. The integrated data included 36,343 cells from 11 patient samples. Cells with more than 50% of reads mapped to mitochondrial genes, those with less than 200 unique genes detected (GEX), those with less than 200 unique peaks detected (ATAC) and those with transcription start site (TSS) enrichment score (as calculated by Signac) less than 1 were removed. After QC filtering the number of cells was reduced to 25,725. Cell type identification was done by using Azimuth algorithm to map the scMultiome GEX data to the scRNA-seq data since they were generated from the same cohort. The labels were then transferred from the scRNA-seq data to the single-cell Multiome data.

Differential gene expression analysis: Differential gene expression testing was done on the log-normalized counts using Seurat’s FindMarkers function with default parameters unless specified otherwise. The statistical test applied was Wilcoxon Rank Sum test with p-values adjusted using Bonferroni correction based on the total number of genes in the dataset. Statistically significant differentially expressed genes were selected by keeping only genes that fall below the adjusted p-value threshold of 0.05. Differential gene expression testing comparing any two conditions (e.g., COVID-19^+^ / *DNMT3A*^MT^ versus COVID-19^+^ / *TET2*^MT^) was always done for each cell type independently (although shown together in the volcano plots for efficient visualization), unless specified otherwise. To make sure that the results were not driven by a single patient sample we applied a leave-one-out approach on all tests where we removed cells from one sample at a time redoing the tests and keeping only the genes that passed the adjusted p-value threshold in all tests. Pathway analysis was done using the Ingenuity Pathway Analysis platform on differentially expressed genes.

Differential DNA accessibility and motif analysis: The differentially accessible peaks were identified by comparing the TF-IDF normalized cut-site counts of any two pair of cell populations using Seurat’s FindMarkers function. Here, the method used was the logistic regression framework along with testing for the number of fragments in peaks as a latent variable. Motif enrichments in peaks were estimated using ChromVAR enrichment scores on the JASPAR2020 motif matrix set.^69, 70^ Enrichment of cut-sites in monocyte and macrophage specific ChIP-seq peaks of select DNA binding proteins were also estimated using ChromVAR by providing ChIP-seq peaks from ReMap2022 as input.^28^ Co-accessibility scores between pairs of peaks were calculated using Cicero v1.3.5.^27^

GoT-ChA^29^: Sequencing reads from the GoT-Cha experiments on two samples (DNMT3A mutants with the R882H mutation) were examined for sufficient sequencing depth. The GoT-Cha experiments yielded 168 million and 151 million reads for samples 1 and 2, respectively. For each sample, we generated four FASTQ files (*_I1_001.fastq.gz, *_R1_001.fastq.gz, *_R2_001.fastq.gz, and *_R3_001.fastq.gz). To precisely locate the cell barcode and mutation site, we randomly selected 5M reads from each FASTQ file and utilized the “sc_seqLogo.py” script from our RSeQC package to generate sequence logos.^71^ Our analysis revealed that the “I1” FASTQ file contains an 8-nt sample barcode (**Supp. Fig. 6B**), while the “R1” FASTQ file contains the 51-nt targeted DNA sequences, with the last three nucleotides representing the genotype of the R882 codon (**Supp. Fig. 6C**). Notably, the codon is CGC (Arginine) for the wildtype and CAC (Histidine) for the mutant, given that the DNMT3A gene is located on the reverse strand. We confirmed that reads from the “R1” file could uniquely map to the targeted DNMT3A locus (chr2:25234324-25234374) on the human reference genome GRCh38/hg38. Furthermore, the “R2” FASTQ file contains the 16-nt cell barcode (**Supp. Fig. 6D**), while the sequences from the “R3” FASTQ files are unknown (**Supp. Fig. 6E**). We then combined the “R1” and “R2” FASTQ files into a single FASTA file, using the cell barcodes as the DNA sequence identifiers, thereby explicitly linking the cell barcode to the DNA sequence (**Supp. Fig. 7**). Any DNA sequence with Phred-scaled quality scores < 30 at the mutation site will be discarded. This preprocessing step not only significantly reduces the file size but also enhances computation speed. For read-level genotyping we used the following method. We utilized an IUPAC (International Union of Pure and Applied Chemistry) string to represent both wildtype and mutation genotypes. In this study, “C” and “T” are denoted as “Y” (**Supp. Fig. 6C**). Then, we employed the motility C++ library (https://github.com/ctb/motility) to match this IUPAC string to each read. Utilizing IUPAC instead of a PWM (Position Weight Matrix) could dramatically enhance computation speed since the differences between wildtype and mutant reads are minimal (only 1 nucleotide change). We found that allowing for 1-mismatch could rescue an additional 10 to 12 million reads (∼7%) in each sample, as compared exact match. Consequently, 77.5% and 75.6% of reads were successfully genotyped for the two samples, respectively. To distinguish real cells from background noise, we employed a methodology similar to CellRanger’s approach in single-cell RNA sequencing analysis. After segregating reads based on cell barcode, we conducted nonparametric kernel density estimation using the “gaussian” kernel function (**Supp. Fig. 8A, B**). The cell barcode density plot revealed three distinct modes likely corresponding to “cell”, “cell-free DNA”, and “empty droplets”, respectively. To determine the cutoff point for cell calling, we identified the local minima (i.e., the changing point where the curve bends) from the density plot. For instance, in sample 1, the local minima for the cell mode is 3.0712, corresponding to 10^3.0712 = 1178 reads. This suggests that a cell with fewer than 1178 reads will be categorized as background (**Supp. Fig. 8A**). A comparison with the Knee plot (or barcode rank plot) demonstrates that the detected threshold aligns well with the elbow point (**Supp. Fig. 8A, B**). As a result, we identified 15185 and 5679 valid cells from samples 1 and 2, respectively. For cell-level genotyping we used the following method. We first calculated the mutant allele fraction (MAF) for each cell using formula described in code deposited on GitHub. Cells with MAF ≤ 0.2 were categorized as homozygous wildtype (WT/WT), cells with MAF ≥ 0.8 were classified as homozygous mutants (Mut/Mut), likely due to Loss of Heterozygosity (LOH), and the remaining cells were designated as heterozygotes (WT/Mut). 40% and 38% of cells in Samples 1 and 2, respectively, were classified as WT/WT (**Supp. Fig. 8C**). The scATAC-seq data from the two samples were analyzed using the cellranger-atac workflow (version 2.1.0), using the reference file (refdata-cellranger-arc-GRCh38-2020-A-2.0.0) downloaded from https://support.10xgenomics.com/single-cell-atac/software/downloads/latest. 8858 and 6255 high-quality cells were identified in Samples 1 and 2, respectively.

## Availability of data and materials

The bulk sequencing, scDNA-seq + proteogenomics, DNA Methylation, scRNA-seq, Multiome (GEX + ATAC) and GoTChA datasets generated and analyzed in this study have been deposited into the NCBI Gene Expression Omnibus (GEO) data base (https://www.ncbi.nlm.nih.gov/geo/) with accession number GSE210435 (reviewers access token: qfcxaoecnjijxct). O-link data has been provided as Supplementary Data. Code and scripts used for analysis are made available in the GitHub repository https://github.com/LabFunEpi/CC_multiomics.

## Notes

### Competing Interest Statement

Dr. Mrinal Patanik has received research funding from Kura Oncology, Epigenetix, Solutherapeutics, Polaris and StemLine Pharmaceuticals.

### Summary of Updates

The revised manuscript reflects an updated and more comprehensive analysis of the data.

## REFERENCES

1. Yang L, Rau R, Goodell MA. DNMT3A in haematological malignancies. Nature reviews Cancer 15, 152-165 (2015).

2. Jaiswal S, Ebert BL. Clonal hematopoiesis in human aging and disease. Science 366, (2019).

3. Jaiswal S, et al. Age-related clonal hematopoiesis associated with adverse outcomes. N Engl J Med 371, 2488-2498 (2014).

4. Jaiswal S, et al. Clonal Hematopoiesis and Risk of Atherosclerotic Cardiovascular Disease. N Engl J Med 377, 111-121 (2017).

5. Buscarlet M, et al. Lineage restriction analyses in CHIP indicate myeloid bias for TET2 and multipotent stem cell origin for DNMT3A. Blood 132, 277-280 (2018).

6. Goyal P, et al. Clinical Characteristics of Covid-19 in New York City. N Engl J Med 382, 2372-2374 (2020).

7. Moore JB, June CH. Cytokine release syndrome in severe COVID-19. Science 368, 473-474 (2020).

8. Onder G, Rezza G, Brusaferro S. Case-Fatality Rate and Characteristics of Patients Dying in Relation to COVID-19 in Italy. JAMA 323, 1775-1776 (2020).

9. Sah P, et al. Asymptomatic SARS-CoV-2 infection: A systematic review and meta-analysis. Proc Natl Acad Sci U S A 118, (2021).

10. Bolton KL, et al. Clonal hematopoiesis is associated with risk of severe Covid-19. Nat Commun 12, 5975 (2021).

11. Zhou Y, et al. Clonal hematopoiesis is not significantly associated with COVID-19 disease severity. Blood 140, 1650-1655 (2022).

12. Duployez N, et al. Clinico-Biological Features and Clonal Hematopoiesis in Patients with Severe COVID-19. Cancers (Basel) 12, (2020).

13. Hameister E, et al. Clonal Hematopoiesis in Hospitalized Elderly Patients With COVID-19. Hemasphere 4, e453 (2020).

14. Zekavat SM, et al. Hematopoietic mosaic chromosomal alterations increase the risk for diverse types of infection. Nat Med 27, 1012-1024 (2021).

15. Netea MG, et al. IL-32 synergizes with nucleotide oligomerization domain (NOD) 1 and NOD2 ligands for IL-1beta and IL-6 production through a caspase 1-dependent mechanism. Proc Natl Acad Sci U S A 102, 16309-16314 (2005).

16. Patnaik MM, et al. DNMT3A mutations are associated with inferior overall and leukemia-free survival in chronic myelomonocytic leukemia. Am J Hematol 92, 56-61 (2017).

17. National Cancer Institutes. Common Terminology Criteria for Adverse Events (CTCAE) v5.0.) (2017).

18. World Health Organization. Clinical management of COVID-19: Living guideline.) (2020).

19. Tulstrup M, et al. TET2 mutations are associated with hypermethylation at key regulatory enhancers in normal and malignant hematopoiesis. Nat Commun 12, 6061 (2021).

20. Encode Project Consortium. An integrated encyclopedia of DNA elements in the human genome. Nature 489, 57-74 (2012).

21. Consortium EP. An integrated encyclopedia of DNA elements in the human genome. Nature 489, 57-74 (2012).

22. Coltro G, et al. Clinical, molecular, and prognostic correlates of number, type, and functional localization of TET2 mutations in chronic myelomonocytic leukemia (CMML)-a study of 1084 patients. Leukemia, (2019).

23. Miles LA, et al. Single-cell mutation analysis of clonal evolution in myeloid malignancies. Nature 587, 477-482 (2020).

24. Stephenson E, et al. Single-cell multi-omics analysis of the immune response in COVID-19. Nat Med 27, 904-916 (2021).

25. Silvin A, et al. Elevated Calprotectin and Abnormal Myeloid Cell Subsets Discriminate Severe from Mild COVID-19. Cell 182, 1401-1418 e1418 (2020).

26. Sandoval L, et al. Characterization and Optimization of Multiomic Single-Cell Epigenomic Profiling. Genes (Basel) 14, (2023).

27. Pliner HA, et al. Cicero Predicts cis-Regulatory DNA Interactions from Single-Cell Chromatin Accessibility Data. Mol Cell 71, 858-871 e858 (2018).

28. Hammal F, de Langen P, Bergon A, Lopez F, Ballester B. ReMap 2022: a database of Human, Mouse, Drosophila and Arabidopsis regulatory regions from an integrative analysis of DNA-binding sequencing experiments. Nucleic Acids Res 50, D316-D325 (2022).

29. Myers RM, Izzo, F, Prieto, T, Mimitou, E, Raviram, R, Chaligne, R, Hoffman, R, Stahl, m, Abdel-Wahab, O, Marcellino, B, Smibert, P, Landau, D. High Throughput Single-Cell Simultaneous Genotyping and Chromatin Accessibility Reveals Genotype to Phenotype Relationship in Human Myeloproliferation. Blood 138, 678 (2021).

30. Mitchell E, et al. Clonal dynamics of haematopoiesis across the human lifespan. Nature 606, 343-350 (2022).

31. Kusne Y, Xie Z, Patnaik MM. Clonal hematopoiesis: Molecular and clinical implications. Leuk Res 113, 106787 (2022).

32. Sano S, Oshima K, Wang Y, Katanasaka Y, Sano M, Walsh K. CRISPR-Mediated Gene Editing to Assess the Roles of Tet2 and Dnmt3a in Clonal Hematopoiesis and Cardiovascular Disease. Circ Res 123, 335-341 (2018).

33. Huang YH, et al. Systematic Profiling of DNMT3A Variants Reveals Protein Instability Mediated by the DCAF8 E3 Ubiquitin Ligase Adaptor. Cancer Discov 12, 220-235 (2022).

34. Nam AS, et al. Single-cell multi-omics of human clonal hematopoiesis reveals that DNMT3A R882 mutations perturb early progenitor states through selective hypomethylation. Nat Genet 54, 1514-1526 (2022).

35. Yamazaki J, et al. Effects of TET2 mutations on DNA methylation in chronic myelomonocytic leukemia. Epigenetics : official journal of the DNA Methylation Society 7, 201-207 (2012).

36. Zhang Q, et al. Tet2 is required to resolve inflammation by recruiting Hdac2 to specifically repress IL-6. Nature 525, 389- 393 (2015).

37. Lasho T, et al. Single cell proteogenomic analysis of aberrant monocytosis in TET2 mutant premalignant and malignant hematopoiesis. Leukemia 37, 1384-1387 (2023).

38. Melenotte C, et al. Immune responses during COVID-19 infection. Oncoimmunology 9, 1807836 (2020).

39. Deshmane SL, Kremlev S, Amini S, Sawaya BE. Monocyte chemoattractant protein-1 (MCP-1): an overview. J Interferon Cytokine Res 29, 313-326 (2009).

40. Chen Y, et al. IP-10 and MCP-1 as biomarkers associated with disease severity of COVID-19. Mol Med 26, 97 (2020).

41. Selimoglu-Buet D, et al. A miR-150/TET3 pathway regulates the generation of mouse and human non-classical monocyte subset. Nat Commun 9, 5455 (2018).

42. Selimoglu-Buet D, et al. Accumulation of classical monocytes defines a subgroup of MDS that frequently evolves into CMML. Blood 130, 832-835 (2017).

43. Patnaik MM, Tefferi A. Chronic myelomonocytic leukemia: 2024 update on diagnosis, risk stratification and management. Am J Hematol, (2024).

44. Kim SH, Han SY, Azam T, Yoon DY, Dinarello CA. Interleukin-32: a cytokine and inducer of TNFalpha. Immunity 22, 131- 142 (2005).

45. Arber DA, et al. The 2016 revision to the World Health Organization classification of myeloid neoplasms and acute leukemia. Blood 127, 2391-2405 (2016).

46. McKenna A, et al. The Genome Analysis Toolkit: a MapReduce framework for analyzing next-generation DNA sequencing data. Genome Res 20, 1297-1303 (2010).

47. Larson DE, et al. SomaticSniper: identification of somatic point mutations in whole genome sequencing data. Bioinformatics 28, 311-317 (2012).

48. Cingolani P, et al. A program for annotating and predicting the effects of single nucleotide polymorphisms, SnpEff: SNPs in the genome of Drosophila melanogaster strain w1118; iso-2; iso-3. Fly (Austin) 6, 80-92 (2012).

49. Fiume M, Williams V, Brook A, Brudno M. Savant: genome browser for high-throughput sequencing data. Bioinformatics 26, 1938-1944 (2010).

50. Landrum MJ, et al. ClinVar: public archive of relationships among sequence variation and human phenotype. Nucleic Acids Res 42, D980-985 (2014).

51. Liu X, Jian X, Boerwinkle E. dbNSFP: a lightweight database of human nonsynonymous SNPs and their functional predictions. Hum Mutat 32, 894-899 (2011).

52. Stenson PD, et al. Human Gene Mutation Database (HGMD): 2003 update. Hum Mutat 21, 577-581 (2003).

53. Tryggvadottir L, et al. Prostate cancer progression and survival in BRCA2 mutation carriers. J Natl Cancer Inst 99, 929- 935 (2007).

54. Robinson D, et al. Integrative clinical genomics of advanced prostate cancer. Cell 161, 1215-1228 (2015).

55. Robinson JT, et al. Integrative genomics viewer. Nat Biotechnol 29, 24-26 (2011).

56. Mule MP, Martins AJ, Tsang JS. Normalizing and denoising protein expression data from droplet-based single cell profiling. Nat Commun 13, 2099 (2022).

57. Stuart T, et al. Comprehensive Integration of Single-Cell Data. Cell 177, 1888-1902 e1821 (2019).

58. Aryee MJ, et al. Minfi: a flexible and comprehensive Bioconductor package for the analysis of Infinium DNA methylation microarrays. Bioinformatics 30, 1363-1369 (2014).

59. Maksimovic J, Gordon L, Oshlack A. SWAN: Subset-quantile within array normalization for illumina infinium HumanMethylation450 BeadChips. Genome Biol 13, R44 (2012).

60. Bernstein BE, et al. The NIH Roadmap Epigenomics Mapping Consortium. Nat Biotechnol 28, 1045-1048 (2010).

61. McLean CY, et al. GREAT improves functional interpretation of cis-regulatory regions. Nat Biotechnol 28, 495-501 (2010).

62. Garapati K, et al. Multiomics single timepoint measurements to predict severe COVID-19 -Authors’ reply. Lancet Digit Health 5, e57 (2023).

63. Zheng GX, et al. Massively parallel digital transcriptional profiling of single cells. Nat Commun 8, 14049 (2017).

64. Wolock SL, Lopez R, Klein AM. Scrublet: Computational Identification of Cell Doublets in Single-Cell Transcriptomic Data. Cell Syst 8, 281-291 e289 (2019).

65. Hao Y, et al. Integrated analysis of multimodal single-cell data. Cell 184, 3573-3587 e3529 (2021).

66. Aran D, et al. Reference-based analysis of lung single-cell sequencing reveals a transitional profibrotic macrophage. Nat Immunol 20, 163-172 (2019).

67. Monaco G, et al. RNA-Seq Signatures Normalized by mRNA Abundance Allow Absolute Deconvolution of Human Immune Cell Types. Cell Rep 26, 1627-1640 e1627 (2019).

68. Stuart T, Srivastava A, Madad S, Lareau CA, Satija R. Single-cell chromatin state analysis with Signac. Nat Methods 18, 1333-1341 (2021).

69. Castro-Mondragon JA, et al. JASPAR 2022: the 9th release of the open-access database of transcription factor binding profiles. Nucleic Acids Res 50, D165-D173 (2022).

70. Schep AN, Wu B, Buenrostro JD, Greenleaf WJ. chromVAR: inferring transcription-factor-associated accessibility from single-cell epigenomic data. Nat Methods 14, 975-978 (2017).

71. Wang L, Wang S, Li W. RSeQC: quality control of RNA-seq experiments. Bioinformatics 28, 2184-2185 (2012).

