## Supplementary Tables for "Single cell multiomic analyses reveal divergent effects of *DNMT3A* and *TET2* mutant clonal hematopoiesis in inflammatory response"

**Supplementary table 1:** Cohort Characteristics of 243 community dwelling patients with COVID-19 stratified by the presence or absence of clonal hematopoiesis (CH).

| **Characteristic/measure** | **All subjects**  **N=243** | **No CH**  **N=171** | **Presence of CH**  **N=72** | **p-value*** |
| --- | --- | --- | --- | --- |
| ***Age at CH eval (years)***  Median (range)  Mean ± sd  <75 years  75+ years  <40 years  40-74 years  75+ years  <60 years  60-74 years  75+ years | 60 (19 – 99)  58.5 ±17.5  201  42  38  163  42  116  85  42 | 57 (19 – 93)  54.3 ±17.0  154  17  37  117  17  93  61  17 | 68.5 (34 – 99)  68.6 ±14.1  47  25  1  46  25  23  24  25 | <0.0001  <0.0001  <0.0001  <0.0001 |
| ***Sex***  Female  Male | 109  134 | 78  93 | 31  41 | 0.82 |
| ***Race/ethnicity group.***  Non-Hispanic White  Black/African American  Asian  Hispanic or Latino  Unknown/missing | 178  23  7  19  16 | 119  16  7  14  15 | 59  7  0  5  1 | 0.071 |
| ***Comorbidities***  Diabetes  Obesity  Hypertension  Coronary heart disease  COPD  More than one  None  Missing  No comorbidities reported  One comorbidity reported  2+ comorbidities reported  Missing | 8  10  40  2  9  79  53  42  53  69  79  42 | 4  10  24  1  5  58  45  24  45  44  58  24 | 4  0  16  1  4  21  8  18  8  25  21  18 | 0.008**  0.033 |

*p-value reflects the comparison of the measures or factors between those with vs. without CH. For continuous measures, a Wilcoxon rank sum test was used, for categorical measures, a chi-square or Fisher’s exact test were used, depending on numbers in each subset.

**based on simulated Fisher’s exact p-value, excluding those with missing comorbidity reporting

Key: COPD: Chronic Obstructive Pulmonary Disease

**Supplementary table 2:** Clinically measured laboratory values in 227 assessable community dwelling patients with COVID-19, stratified by the presence or absence of clonal hematopoiesis (CH)

| **Characteristic/measure**  Median (range)  Mean  *# missing* | **All subjects**  **N=227** | **No CH**  **N=171** | **Presence of CH**  **N=56** | **p-value*** |
| --- | --- | --- | --- | --- |
| Hemoglobin | 13.35 (7.1 – 18.6)  13.02  *83* | 13.6 (7.7 – 18.6)  13.19  *67* | 12.55 (7.1 – 16.8)  12.59  *16* | 0.099 |
| Platelets | 191.5 (0.91 – 609)  209.67  *83* | 194.5 (0.91 – 609)  212.78  *67* | 186.5 (69 – 467)  201.6  *16* | 0.41 |
| White blood cell count | 5.9 (0.8 – 25.0)  7.01  *83* | 6.1 (0.8 – 25)  7.21  *67* | 5.6 (1.1 – 16.3)  6.49  *16* | 0.31 |
| Absolute neutrophil count | 4.36 (0.26 – 186)  7.58  *87* | 4.73 (0.26 – 186)  8.13  *70* | 4.08 (0.82 – 60.9)  6.16  *17* | 0.15 |
| Absolute monocyte count | 0.44 (0.03 – 17.3)  0.75  *87* | 0.44 (0.1 – 11.3)  0.67  *70* | 0.44 (0.03 – 17.3)  0.95  *17* | 0.86 |
| Absolute lymphocyte count | 0.89 (0.03 – 21.2)  1.29  *87* | 0.89 (0.1 – 13.9)  1.15  *70* | 0.81 (0.03 – 21.2)  1.67  *17* | 0.76 |
| C-reactive protein | 82.4 (0 – 1240)  118.1  *95* | 92.25 (0 – 1240)  127.74  *77* | 66.2 (11.2 – 400)  94.26  *18* | 0.14 |
| Ferritin | 466 (35 – 4157)  742.3  *103* | 515 (38 – 4157)  772.8  *84* | 456 (35 – 3686)  670.5  *19* | 0.62 |
| IL-6 | 20.4 (1 – 400)  39.07  *125* | 20.4 (1 – 400)  41.04  *101* | 19.9 (1 – 320)  34.75  *24* | 0.81 |

**Supplementary table 3: Cohort cytokine measures for community dwelling patients with COVID-19, as stratified by the presence or absence of clonal hematopoiesis (CH).**

| Characteristic/measure  Median (range)  Mean  *# < lower limit of detection* | All subjects  N=165* | No CH  N=128 | Presence of CH  N=37 | p-value** |
| --- | --- | --- | --- | --- |
| TNF | 5 (5 – 38.5)  7.73  *124* | 5 (5 – 38.5)  7.74  *98* | 5 (5 – 17.4)  7.7  *26* | 0.54 |
| IL-6 | 2.5 (2.5 – 320)  23.02  *111* | 2.5 (2.5 – 320)  22.8  *90* | 2.5 (2.5 – 320)  23.8  *21* | 0.20 |
| IFN-b | *160* | *124* | *36* |  |
| IL-10 | *156* | *120* | *36* |  |
| MCP-1 | 238.5 (20 – 4408)  353.6  6 | 209.4 (20 – 4408)  355.6  6 | 274.6 (77.4 – 1249)  346.4  0 | 0.045 |
| IL-1b | *160* | *124* | *36* |  |
| IFN-g | *161* | *124* | *37* |  |
| MIP-1a | *160* | *123* | *37* |  |
| GM-CSF | *164* | *127* | *37* |  |
| sIL-2a | 695.6 (20 – 4100)  935.5  *2* | 669.5 (20 – 3136)  913.3  *2* | 766.7 (235 – 4100)  1012.1  *0* | 0.23 |
| IFN-a | 10 (10 – 221)  21.39  *90* | 10 (10 – 221)  20.35  *72* | 21.6 (10 – 177.6)  24.97  *18* | 0.25 |
| IL-18 | 282.6  (32.5 – 1708)  370.6  *5* | 267.9  (32.5 – 1708)  367.8  *4* | 310.7  (32.5 – 1211)  380.2  *1* | 0.56 |

*Total number of subjects in data base is N=243, but 78 subjects had missing cytokine measures (43 without CH, 35 with CH)

**p-value reflects the comparison of the measures or factors between those with vs. without CH. For continuous measures, a Wilcoxon rank sum test was used, for categorical measures, a chi-square or Fisher’s exact test were used, depending on numbers in each subset.

**Supplementary table 4:** Cohort disease characteristics of community dwelling patients with COVID-19 with regards to clinical presentation and outcomes, stratified by the presence or absence of clonal hematopoiesis (CH).

| **Characteristic/measure** | **All subjects**  **N=243** | **No CH**  **N=171** | **Presence of CH**  **N=72** | **p-value*** |
| --- | --- | --- | --- | --- |
| ***Hospitalization?***  No  Yes | 110  133 | 77  94 | 33  39 | 0.99 |
| ***Oxygen required?***  No  Yes  missing | 42  113  88 | 27  77  67 | 15  36  21 | 0.79 |
| ***Max oxygen requirements***  Invasive mechanical ventilation  Noninvasive ventilation  Missing | 19  93  130 | 12  65  94 | 7  29  36 | 0.68 |
| ***Cytokine release syndrome (CRS)***  No  Yes | 119  124 | 81  90 | 38  34 | 0.53 |
| ***CRS grade by CTCAE.V5.0 criteria***  0 (None)  1 (mild)  2 (moderate)  3 (severe)  4 (life-threatening)  5 (death) | 119  12  55  43  1  13 | 81  12  40  32  1  5 | 38  0  15  11  0  8 | 0.018 |
| ***CRS grade by WHO scale***  1 (Ambulatory, no limits)  2 (Ambulatory, limits)  3 (Hosp., no O2 or med care)  4 (Hosp., no O2, but med care)  5 (Hosp, supp O2)  6 (Hosp, req NIV or HFNC)  7 (Hosp, iMV or ECMO)  8 (Death)  Median (range)  Mean ± sd  Missing | 105  4  1  20  2  83  16  10  4 (1 – 8)  3.7 ± 2.6  2 | 71  3  1  17  2  59  13  3  4 (1 – 8)  3.7 ± 2.5  2 | 34  1  0  3  0  24  3  7  4 (1 – 8)  3.7 ± 2.76  0 | 0.12  0.90 |
| ***Acute Lung Injury***  No  Yes  Missing | 55  85  103 | 40  62  69 | 15  23  34 | 0.61 |
| ***Acute Kidney Injury***  No  Yes  Missing | 112  28  103 | 84  18  69 | 28  10  34 | 0.33 |
| ***Acute respiratory distress syndrome***  No  Yes  Missing | 106  34  103 | 80  22  69 | 26  12  34 | 0.30 |
| ***Multi-organ dysfunction syndrome***  No  Yes  Missing | 131  9  103 | 98  4  69 | 33  5  34 | 0.095 |
| ***Thrombosis***  No  Yes  Missing | 122  17  104 | 88  13  70 | 34  4  34 | 0.62 |
| ***Therapy received for COVID***  None  Remdesivir  Paxlovid  Monoclonal antibody  Convalescent plasma  Clinical trial  More than one | 139  51  1  4  8  3  37 | 100  34  0  4  6  1  26 | 39  17  1  9  2  2  11 | 0.33 |
| ***Status at last f/u***  Alive  Dead | 226  17* | 165  7 | 62  10 |  |

*n=12 patients reported as ‘dead’ in event status, but 2 additional patients had grade 5 events reported and are included in this number
